# Mitochondrial activity directs nutrient uptake to control early human embryonic patterning

**DOI:** 10.64898/2026.05.20.725854

**Authors:** Shiyu Bian, Marta M. Marcheluk, Ana R. Lima, Songyang Li, Christopher J. Price, Veronique Azuara, Ivana Barbaric, Tristan A. Rodriguez

**Affiliations:** National Heart and Lung Institute, Imperial College London, UK; Department of Metabolism, Digestion, and Reproduction, Imperial College London, UK; Department of Biomedical Science, The University of Sheffield, UK

**Keywords:** mitochondrial dysfunction, mtDNA, nutrient uptake, BMP signalling, patterning, human development

## Abstract

How a uniform field of cells responds to inductive cues to form distinct, spatially organised, lineages is a question of much interest. In human gastruloids, a BMP4 signalling gradient directs the self-organization of embryonic stem cells (hESCs) into the three germ-layers. But what regulates differential responses to BMP4 is poorly understood. Here we demonstrate that mitochondrial activity is key, as by directing nutrient uptake it determines the threshold of BMP signalling. We show that decreased mitochondrial activity stimulates fatty acid uptake via macropinocytosis and subsequent β-oxidation and that this increases the BMP4 response. However, we also find that disease causing mitochondrial DNA mutations impair dynamic switching between glucose and fatty acid utilisation, causing hESCs to be unable to spatially organise the mesoderm, endoderm and ectoderm layers in gastruloids. These results suggest that the correct interpretation of morphogen gradients during gastrulation emerges through the crosstalk of signalling inputs and nutrient utilisation.

## Background

Mitochondrial oxidative phosphorylation (OxPhos) generates ∼90% of the body’s energy through ATP synthesis. The catabolism of glucose via glycolysis and pyruvate oxidation, of fatty acids via β-oxidation, or of amino acids via oxidative deamination and transamination, is used by the tricarboxylic acid cycle to generate substrates for the electron transport chain (ETC). The ETC, comprising four protein complexes (Complexes I-IV) embedded in the inner mitochondrial membrane and two mobile electron carriers (ubiquinone and cytochrome c), uses these substrates to generate ATP. In addition to their roles in cellular bioenergetics, mitochondria perform a plethora of other functions, most notably the regulation of apoptosis, the biosynthesis of macromolecules and as a signalling hub^1^.

Mitochondria have their own DNA and in humans, the mitochondrial DNA (mtDNA) encodes 13 components of the OxPhos machinery, 22 mitochondrial tRNAs and 2 rRNA. Due to its proximity to the ETC, mtDNA is susceptible to oxidative damage, leading to a mutation rate that is higher than that of nuclear DNA^2^. Many of these mutations disrupt mitochondrial gene expression or impair OxPhos, thereby compromising cellular bioenergetics. Mutations in these mtDNA genes cause primary mitochondrial disorders, that roughly affect 5-20 in 100,000 of the population and have no effective treatment^3–5^. These can be present from birth to old age, and although clinically heterogeneous, frequently affect the nervous system, as this organ system has one of the highest energy demands^6^. The relative ratio of mutant to wild-type mtDNA copies in a cell (heteroplasmy) determines the degree of disease manifestation, with most pathogenic mtDNA mutations needing to exceed a threshold of 50%–90% to induce OxPhos dysfunction^7^. However, this threshold varies according to the type of mutation and the cell type. Although high pathogenic mtDNA mutations have been associated with early embryo loss^8^, what roles mitochondrial activity plays in embryonic development and how mtDNA mutations impact these roles is not fully understood.

The early post-implantation period of embryonic development and the onset of gastrulation is a major developmental landmark. It involves adapting not only to the new environment provided by the uterus after implantation but also to the changes caused by the specification of the three germ-layers, endoderm, mesoderm and ectoderm. About 60-80% of human conceptions end in a miscarriage, with 30% failing between weeks 2 and 6 of gestation, around the time of gastrulation^9, 10^. Furthermore, many developmental abnormalities are thought to originate during this period of development^11^.

Nutrients are emerging as cues that direct cell fate and behaviour during the early embryonic development. For example, glucose directs mesoderm specification^12–14^ and migration^12^ in mouse and humans, the mode of amino acid uptake defines the pluripotent states^15^ and lipids regulate epiblast morphogenesis in mouse^16^ and endoderm fate in human pluripotent stem cells^17^. Despite this interest, what determines which nutrients are selectively utilised during gastrulation, how they modulate the signalling responses that govern embryonic patterning and how they are affected by disease causing mutations is still poorly understood.

Over the last few years, stem cell-based models of gastrulation or “gastruloids” have emerged as valuable systems to study early human gastrulation^18^. One such model is 2D gastruloid micropatterns, where BMP4 induces human embryonic stem cells (hESCs) to self-organise into an outer endodermal layer, an intermediate mesodermal layer and an inner ectodermal layer^19^. In 2D hESC micropatterns, an exogenous BMP4 signal induces the embryonic germ layers to organise into an outer endodermal layer, an intermediate mesodermal layer and an inner ectodermal layer^19–26^. What determines the differential response to BMP4 is a question of much interest.

Here, we have investigated the impact that mtDNA mutations have on hESC function and patterning. For this we have generated cells carrying mutations in mtND6, a key component of Complex I of the ETC. We find that high heteroplasmy mutations reduce OxPhos and change nutrient dependency from glucose to fatty acid oxidation to sustain their proliferation. In turn, these changes increase the sensitivity of differentiating cells to BMP4 signals and consequently disrupt organization of the 3 germ-layers in 2D micropatterns. Our work therefore provides novel insight into the how mitochondrial activity directs nutrient uptake to regulate the signalling that imparts embryonic patterning and how mtDNA mutations disrupt this process.

## Results

### High heteroplasmy mutations in mtND6 reduce OxPhos in hESCs

Here we aimed to determine the effects that mtDNA mutations have on early embryonic patterning in humans. For this, we first set out to establish mutant hESCs carrying mutations in mtDN6, a key component of Complex I of the ETC and for which mutations have been described in Leigh Syndrome^27–33^, Mitochondrial Encephalopathy, Lactic Acidosis, and Stroke-like episodes (MELAS)^34^ and Leber’s hereditary optic neuropathy (LHON)^35–45^. To introduce mutations, cells were transfected with two plasmids encoding the DddA-derived cytosine base editors (DdCBE)^46^. Each plasmid carries a TALE arm and either a GFP or tdTomato reporter to enable FACS sorting of double-positive cells. Each TALE arm encodes half of the DddA deaminase that bind adjacent sequences flanking the target site, reconstituting the split DddA deaminase upon co-binding. Lines were established from sorted single cells and sequencing revealed the presence of mutations within the targeted regions of mtND6 (Fig.1a and Fig.S1a). We then classified these lines according to their level of heteroplasmy for the site with the highest mutation frequency into high (>75%), medium (40-75%) and low (<40%)(Fig.1b and Fig.S1b). Our analysis also revealed that some lines contained a small number of off-target mtDNA mutations that were present at low heteroplasmy levels (Table S1). Importantly, the level of mtND6 heteroplasmy at individual mutant sites remained stable during repeat passaging of the cells (Fig.1c), suggesting that there was no strong selection bias against the mutations generated.

**Figure 1.**
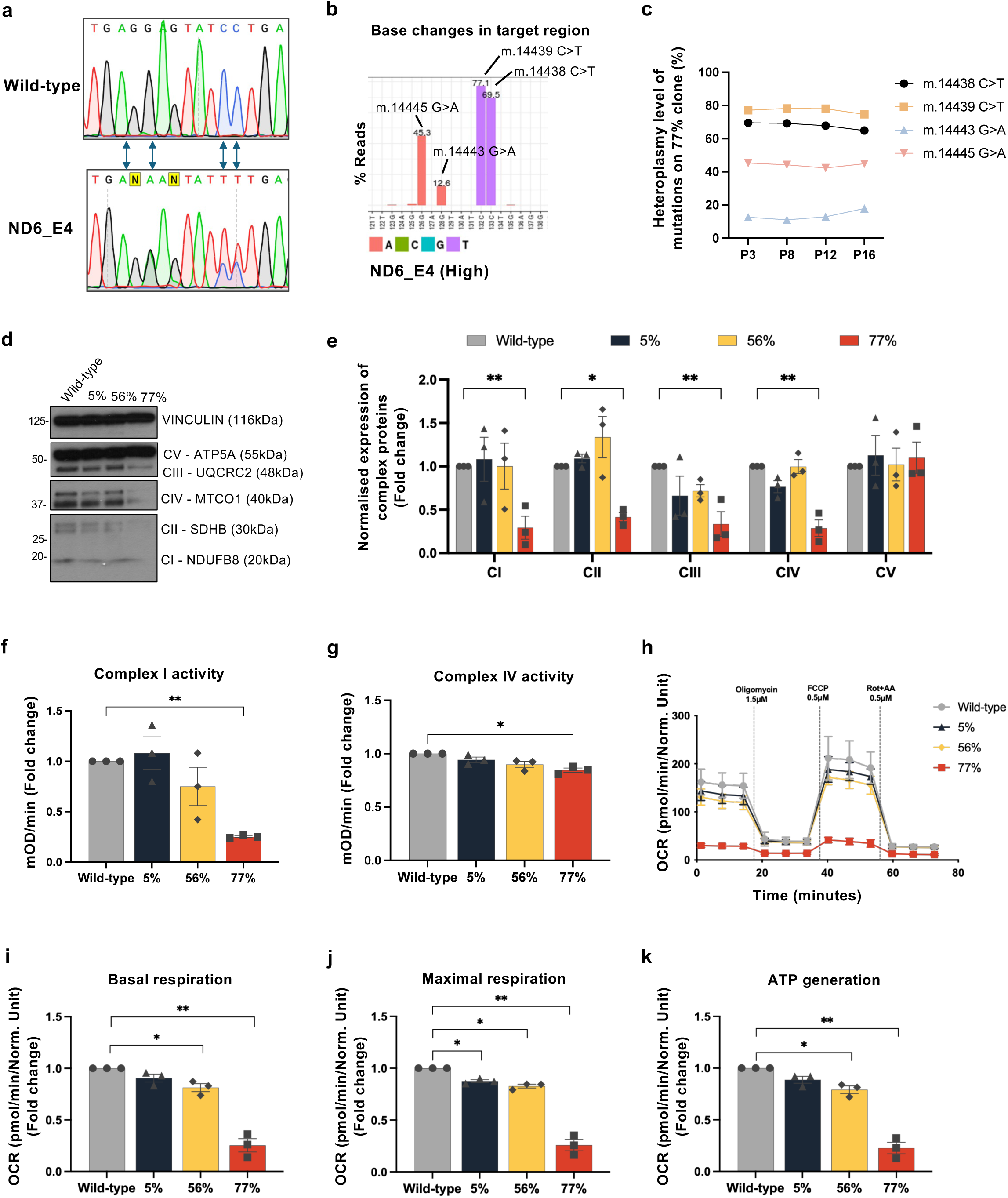
Generation and molecular characterization of mtND6 mutant hESC clones with different heteroplasmy levels. (a) Sanger sequencing chromatograms of the mtND6 target region in wild-type and ND6_E4 hESCs. Arrows indicate edited bases introduced by DdCBE. (b) Heteroplasmy levels of m.14438C>T, m.14439C>T, m.14443G>A, and m.14445G>A, quantified by amplicon sequencing. Values above each bar indicate the corresponding heteroplasmy percentage. (c) Heteroplasmy levels of individual mtDNA mutations in the 77% ND6_E4 clone measured at passages P3, P8, P12 and P16. (d) Immunoblot analysis of OxPhos complex subunits in wild-type cells and mtND6 mutant clones with 5%, 56% and 77% heteroplasmy. VINCULIN was used as a loading control. (e) Quantification of NDUFB8 (Complex I), SDHB (Complex II), UQCRC2 (Complex III), MTCO1 (Complex IV), ATP5A (Complex V), protein levels normalised to wild-type levels. (f-g) Enzymatic activity of mitochondrial Complex I (f) and Complex IV (g), measured by absorbance at 450 nm and 550 nm, respectively. (h) Oxygen consumption rate (OCR) profiles of wild-type and mtND6 mutant hESCs (5%, 56%, and 77%) measured using the Seahorse Mito Stress Assay. (i-k) Quantification of basal respiration (i), maximal respiration (j), and ATP-linked respiration (k), calculated from OCR measurements. FCCP, carbonyl cyanide 4-(trifluoromethoxy)phenylhydrazone. Data are shown as mean ± SEM from n=3 independent biological replicates. Statistical significance was assessed using two-way ANOVA with Šídák’s post-hoc test for (e) and two-tailed one-sample t-test for (f, g and i-k); *p < 0.05, **p < 0.01, ****p<0.0001.

To establish how these mutations affect ETC assembly and function we first measured the expression of key subunits involved in the assembly of the five respiratory complexes. This revealed that high heteroplasmy lines grown in mTeSR^TM^ Plus not only displayed disrupted Complex I assembly but also disruption of Complexes II-IV. In contrast, Complex V remained unaffected (Fig.1d-e). Disruption of the assembly of multiple complexes has been found for other Complex I mutations, presumably due to impaired respiratory supercomplex formation^47, 48^.

Analysis of Complex activity revealed that whilst Complex I function was reduced by about 75% in high load mutant lines, Complex IV only showed a small reduction in activity (Fig.1f-g). Most pathogenic mtDNA mutations need to exceed a threshold of 50%–90% to induce OxPhos dysfunction^7^. Importantly, we found that the high heteroplasmy mtND6 mutations generated compromised respiration rates, causing about a 75% reduction in basal and maximal respiration, as well as a 77.5% reduction in ATP generation through the ETC (Fig.1h-k, Fig.S1c-d). This reduction was specific to high heteroplasmy lines, as those with 56% heteroplasmy or below only showed marginally lower respiration parameters than control cells or no change (Fig.1h-k), even if they also had similar levels of off-target mtDNA mutations (Table S1).

Analysis of the mitochondrial membrane potential, an indication of OxPhos efficiency, revealed no change in high heteroplasmy lines compared to controls (Fig.S1e-f). This is in line with the decreased proton leak observed in these cells (Fig.S1c-d), as a reduced leak would buffer a lower respiration efficiency. These cells also displayed slightly elevated mitochondrial ROS levels (Fig.S1g-h). This agrees with the observed Complex I disruption, as its inhibition would lead to a build-up of electrons in the context of oxygen availability, leading to ROS.

In conclusion, together these results indicate that mtND6 mutations reduce oxidative respiration to a degree that is directly related to their level of heteroplasmy.

### MtND6 mutation induces a switch in nutrient uptake requirements in hESCs

To establish how mtND6 mutations affect hESC fitness we compared the proliferation rate of cells with different heteroplasmy levels. We observed that in mTeSR^TM^ Plus all lines proliferated at a similar rate to controls irrespective of their level of heteroplasmy (Fig.2a-b and Fig.S2a). mTeSR^TM^ Plus is a nutrient rich media that could provide metabolic flexibility allowing cells to rely on compensatory pathways. We tested this possibility by analysing the behaviour of high heteroplasmy mutant cells in the nutrient poor E8 media (Fig.S2b). We observed that mtND6 hESCs grown in E8 displayed severely compromised growth (Fig.2c-d). Furthermore, we found that unlike control cells that displayed a glucose-dependent proliferation in E8, high heteroplasmy mtND6 cells did not show any proliferation increase when glucose was added back to glucose-depleted media (Fig.2e-f). This was not because these cells were unable to perform glycolysis, as glucose uptake and glycolytic rates were similar in mutant cells to control cells (Fig.2g-l). These results suggest that high heteroplasmy mutations in mtND6 change how hESCs utilise their nutrients to support growth.

**Figure 2.**
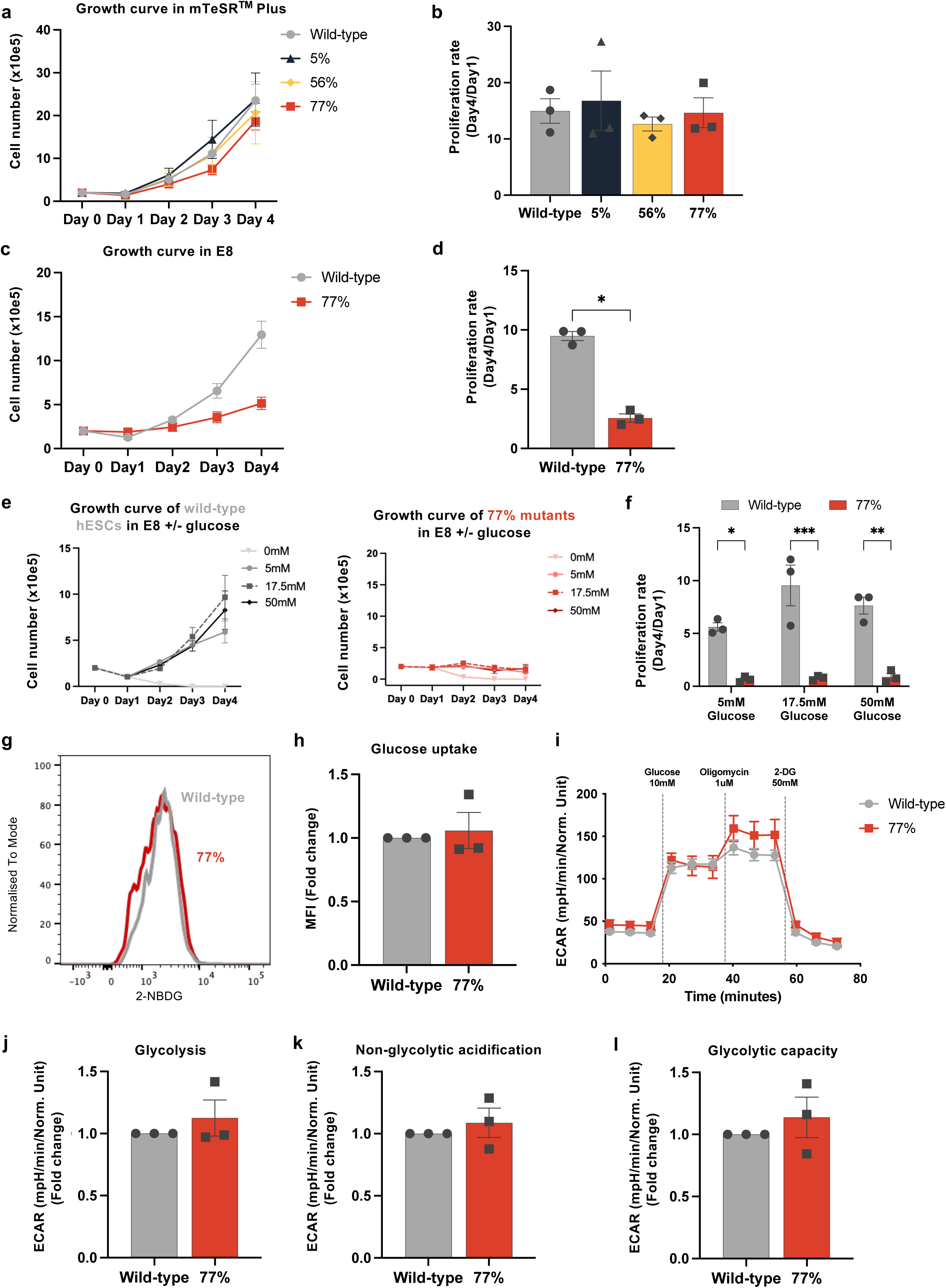
Mitochondrial dysfunction causes a proliferation defect specifically in a nutrient-poor environment. (a) Growth curves of wild-type and mtND6 mutant hESCs with 5%, 56%, and 77% heteroplasmy cultured in mTeSR^TM^ Plus medium. Cell numbers were quantified over 4 days. (b) Proliferation rates of wild-type and mtND6 mutant hESCs (5%, 56%, and 77%) in mTeSR^TM^ Plus, calculated from growth curves in (a). (c) Growth curves of wild-type and 77% mtND6 mutant hESCs cultured in E8 medium. (d) Proliferation rates of wild-type and 77% mtND6 mutant hESCs in E8, calculated from growth curves in (c). (e) Growth curves of wild-type (left) and 77% mtND6 mutant hESCs (right) cultured in E8 supplemented with increasing concentrations of glucose (0, 5, 17.5, and 50 mM). (f) Proliferation rates of wild-type and 77% mtND6 mutant hESCs in E8 supplemented with different glucose concentrations. (g) Representative flow cytometry histograms of 2-(N-(7-Nitrobenz-2-oxa-1,3-diazol-4-yl)Amino)-2-Deoxyglucose (2-NBDG) fluorescence in wild-type and 77% mtND6 mutant hESCs cultured in mTeSR^TM^ Plus. (h) Quantification of 2-NBDG median fluorescence intensity (MFI) in wild-type and 77% mtND6 mutant hESCs, expressed as fold change relative to wild-type. (i) Extracellular acidification rate (ECAR) profiles of wild-type and 77% mtND6 mutant hESCs cultured in mTeSR^TM^ Plus, measured using the Seahorse Glycolysis Stress Test, with sequential addition of glucose (10 mM), oligomycin (1 μM), and 2-deoxyglucose (2-DG; 50 mM). (j-l) Quantification of glycolysis (j), non-glycolytic acidification (k) and glycolytic capacity (l) derived from ECAR measurements and expressed as fold change relative to wild-type. Data are shown as mean ± SEM from n=3 independent biological replicates. Statistical significance was assessed using one-way ANOVA with Geisser-Greenhouse correction and Dunnett’s multiple comparisons test for (b), or two-tailed paired t-test for (d), or two-way ANOVA with Tukey’s post-hoc test for (f) and two-tailed one-sample t-test for (h and j-l); *p < 0.05, **p < 0.01, ***p<0.001, ****p<0.0001.

Our observation that mutant cells proliferated normally in mTeSR^TM^ Plus but not E8 led us to hypothesize that these cells are reliant for their growth on a nutrient present in mTeSR^TM^ Plus but not in E8. We therefore compared the formulations of both media (Fig.S2b) and sequentially tested individual nutrients. When this was done, we found that in E8, bovine serum albumin (BSA) was the only component that was sufficient to increase to control levels the proliferation rate of high heteroplasmy mtND6 cells (Fig.3a-b and Fig.S3a-d). This indicates that mtND6 mutation induces hESCs to become dependent on albumin uptake for their growth.

**Figure 3.**
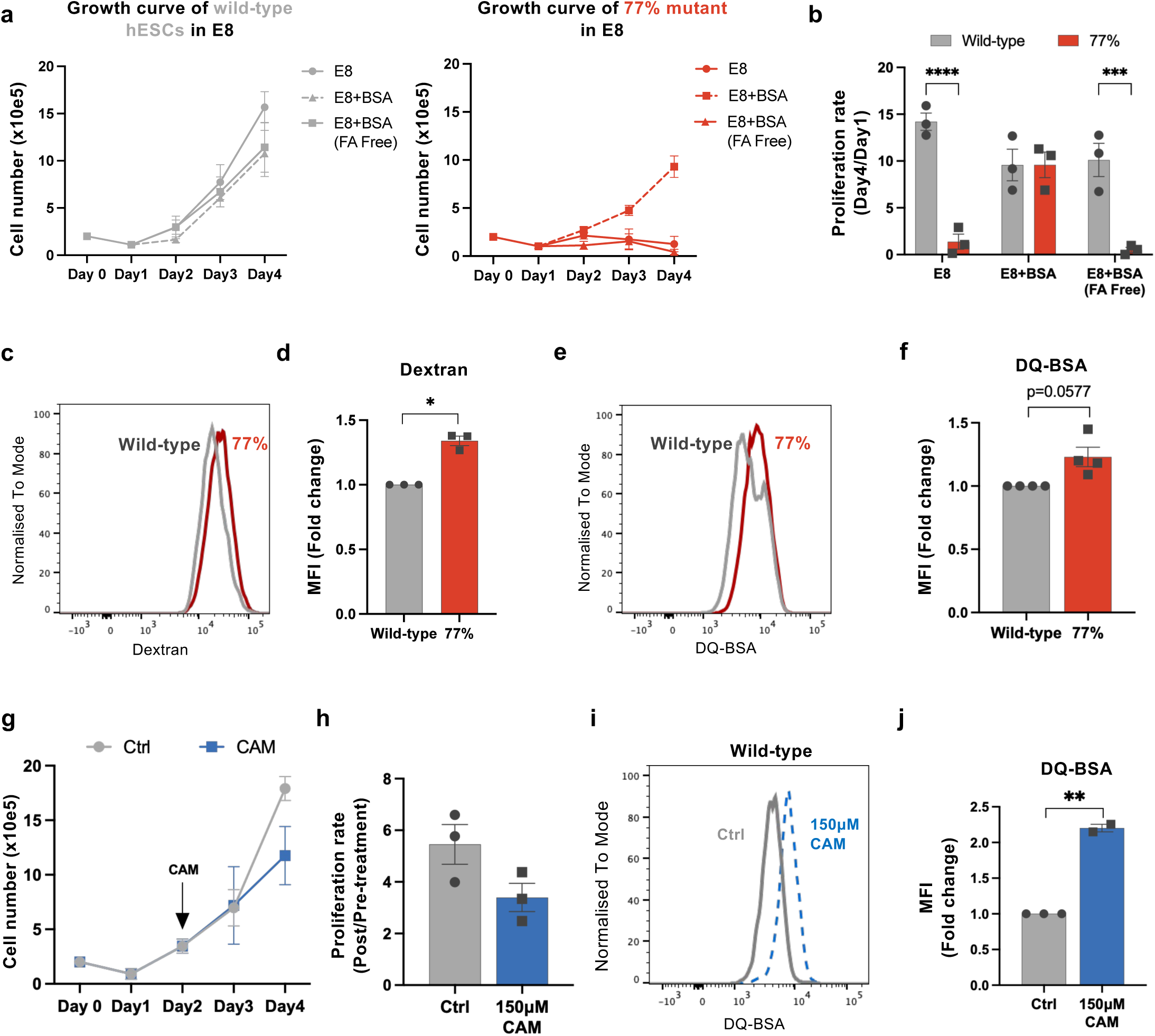
mtND6 mutation causes metabolic rewiring and enhances macropinocytosis in hESCs. (a) Growth curves of wild-type (left) and 77% mtND6 mutant hESCs (right) cultured in E8, E8 supplemented with BSA, or E8 supplemented with fatty acid-free BSA. (b) Proliferation rates of wild-type and 77% mtND6 mutant hESCs under the conditions shown in (a). (c) Flow cytometry histograms of fluorescent dextran uptake in wild-type (grey) and 77% mtND6 mutant hESCs (red) cultured in mTeSR^TM^ Plus medium. (d) Median fluorescence intensity (MFI) of dextran uptake in wild-type and 77% mtND6 mutant hESCs in E8, expressed as fold change relative to wild-type. (e) Flow cytometry histograms of DQ-Red BSA uptake in wild-type (grey) and 77% mtND6 mutant hESCs (red) cultured in mTeSR^TM^ Plus medium. (f) MFI of DQ-Red BSA uptake in wild-type and 77% mtND6 mutant hESCs in E8, expressed as fold change relative to wild-type. (g) Growth curves of wild-type hESCs treated with chloramphenicol (CAM) or vehicle control. (h) Proliferation rates of wild-type hESCs following 150μM CAM treatment, calculated as the ratio of post-treatment to pre-treatment proliferation rates (over 48 hours). (i) Flow cytometry histograms of DQ-Red BSA uptake in wild-type hESCs under control conditions (Ctrl) or following 48 hours 150μM CAM treatment. (j) Quantification of DQ-Red BSA uptake in wild-type hESCs under control conditions or following CAM treatment, expressed as fold change relative to control. Data are shown as mean ± SEM from n=3 or 4 independent biological replicates. Statistical significance was assessed using two-way ANOVA with Tukey’s post-hoc test for (b), or two-tailed one-sample t-test for (d, f and j) and two-tailed paired t-test for (h); *p < 0.05, **p < 0.01, ***p < 0.001, ****p < 0.0001.

Macropinocytosis, or the non-selective uptake of nutrients from the environment, is an important mechanism for albumin uptake in cells^49^, and mouse ESCs rely on it to ensure an adequate supply of amino acids^15^. This led us to investigate if mutant cells exhibited differences in macropinocytosis when compared to controls. Interestingly, we observed that in mTeSR^TM^ Plus media, high heteroplasmy cells showed higher uptake than control cells of both dextran and DQ-BSA, two substrates taken up by macropinocytosis (Fig.3c-f). Furthermore, we found that inducing mitochondrial dysfunction in wild-type hESCs with a dose of chloramphenicol that partially inhibits mitochondrial translation (Fig.S4a-b) and causes only a small proliferative impairment (Fig.3g-h), stimulates macropinocytosis (Fig.3i-j). These findings suggest that mitochondrial dysfunction induces a switch in nutrient utilisation involving the uptake of albumin via macropinocytosis.

### Mutant mtND6 hESCs increase fatty acid uptake to compensate for their OxPhos deficit

Albumin is the main carrier of fatty acids in the blood^50^, but can also be used as source of amino acids^51^. For this reason, we set out to distinguish if the proliferation increase induced by BSA was due to the import of fatty acids or protein. Importantly, we observed that neither fatty acid free BSA or a lipid concentrate alone had any effect on the proliferation of mutant cells (Fig.3a-b and Fig.S3a-b). This indicates that high heteroplasmy mtND6 mutant hESCs import fatty acids to sustain their growth and that these fatty acids need to be coupled to albumin to enter the cell.

To further probe the possibility that mtND6 mutations induce a change in nutrient utilisation to fatty acid oxidation we compared the transcriptional profile of control and mutant cells in mTeSR^TM^ Plus and E8 media. In both conditions we observed the upregulated pathways to be primarily related to lipid metabolism, including cholesterol and sterol biosynthesis and metabolism, as well as fatty acid response (Fig.4a-d and Table S2a-b). These changes were accompanied by an increase in the expression of *ACLY* (ATP citrate lyase) and *ACSS2* (acyl-CoA synthetase short-chain family member 2), two enzymes required for the generation of cytosolic acetyl-CoA, a precursor for both fatty acid and cholesterol biosynthesis (Fig.4a and c and e-f). To test the functional significance of these changes we analysed the effect of SB-204990 and ETC-1002 (ACLY inhibitors), of ACSS2i (an ACSS2 inhibitor) and of pravastatin (a statin-class drug that inhibits HMG-CoA reductase, suppressing cholesterol synthesis). We found that none of these inhibitors had a differential effect on high heteroplasmy mtND6 mutant versus control cell (Fig.S3e-l). This supports the hypothesis that fatty acid uptake from the environment rather than an increase in fatty acid and cholesterol biosynthesis is what sustains proliferation upon mtND6 mutation.

**Figure 4.**
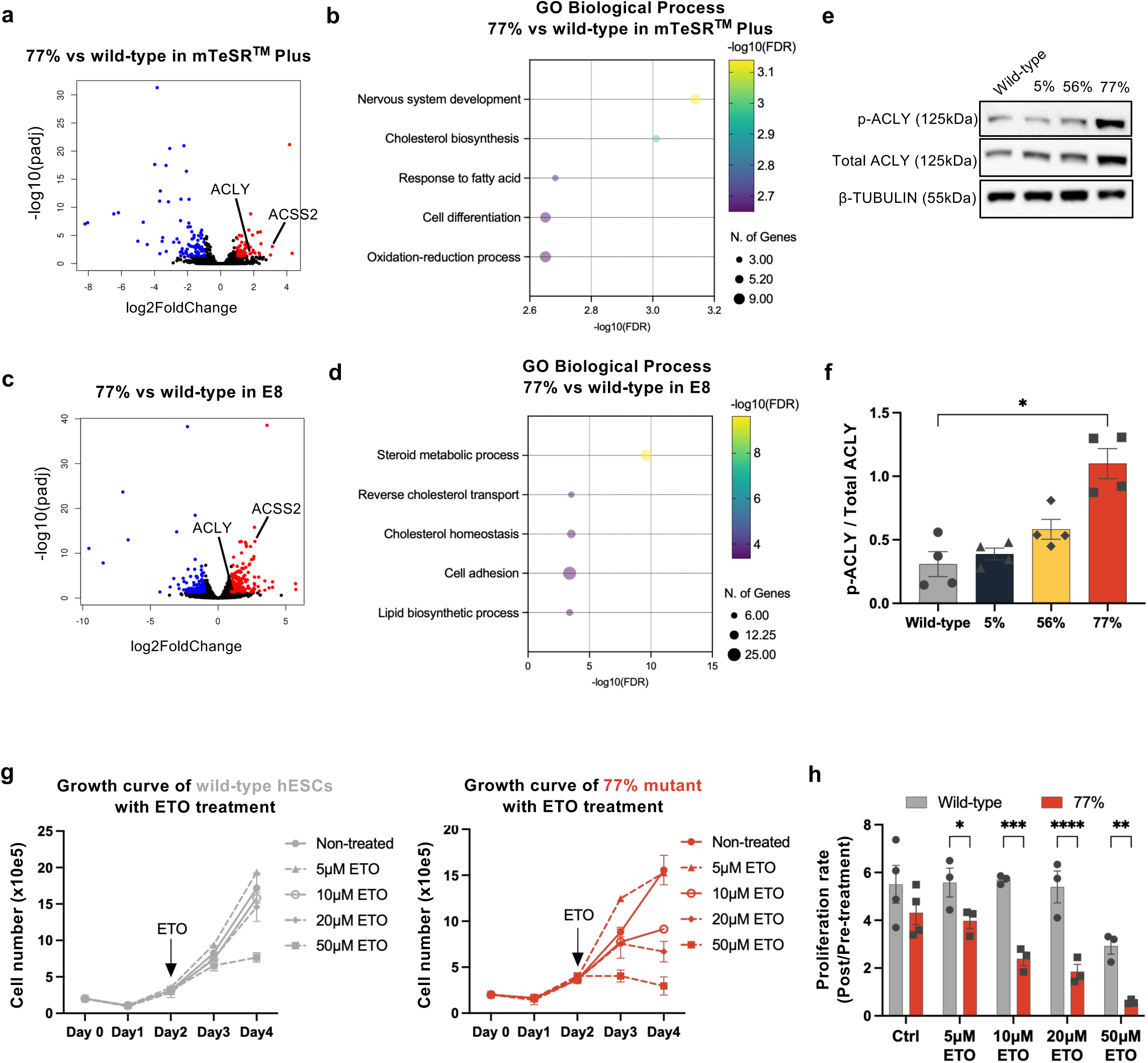
Transcriptomic and functional evidence for altered fatty acid metabolism in mtND6 mutant hESCs. (a and c) Volcano plot of differentially expressed genes (DEGs) between 77% mtND6 mutant and wild-type hESCs cultured in mTeSR^TM^ Plus medium (a) or E8 medium (c). (b and d) Bubble plot of Gene Ontology (GO) biological process enrichment analysis of DEGs identified in 77% mtND6 mutant versus wild-type hESCs cultured in mTeSR^TM^ Plus medium (b) or E8 medium (d), with bubble size representing the number of genes associated with each term and colour indicating - log10(FDR). FDR, false discovery rate. (e) Representative immunoblot analysis of phosphorylated ACLY (p-ACLY), total ACLY and β-TUBULIN (loading control) in wild-type 5%, 56%, and 77% mtND6 mutant hESCs. (f) Quantification of the p-ACLY/Total-ACLY ratio and expressed as fold change relative to wild-type hESCs. (g) Growth curves of wild-type (left) and 77% mtND6 mutant (right) hESCs treated with 5-50 μM etomoxir (ETO) or vehicle control, with the time point of etomoxir addition indicated by the arrow. (h) Proliferation rates of wild-type and 77% mtND6 mutant hESCs following Etomoxir (ETO) or vehicle control treatment, calculated as the ratio of post-treatment to pre-treatment proliferation rates (48 hours). Data are shown as mean ± SEM from n=3 or 4 independent biological replicates. Statistical significance was assessed using one-way ANOVA with Tukey’s post-hoc test for (f), or two-way ANOVA with Tukey’s post-hoc test for (h); *p < 0.05, **p < 0.01, ***p < 0.001, ****p < 0.0001.

We next tested the possibility that mutant cells were relying on fatty acid oxidation rather than glucose catabolism for their proliferation. For this we treated cells with etomoxir, a carnitine palmitoyltransferase-1 (CPT-1) inhibitor, that prevents the transport of fatty acyl chains into the mitochondria for their subsequent oxidation during OxPhos. We observed that after 48 hours treatment, whilst the proliferation of wild-type cells was unaffected by up to 20μM etomoxir, high heteroplasmy mtND6 cells displayed impaired proliferation with 10 and 20μM etomoxir (Fig.4g-h). Together, these results indicate that mtDNA mutation in mtND6 causes hESCs to increase their reliance on the uptake and oxidation of fatty acids, instead of glucose oxidation.

### Mitochondrial dysfunction disrupts pattern formation in 2D gastruloids

We next wanted to understand how the mitochondrial dysfunction induced by mtND6 mutations impact the pluripotent status of hESCs, their ability to differentiate into the three germ-layers and undergo pattern formation. Analysis of the protein expression of pluripotency markers OCT4 (POU5F1) and NANOG indicated no difference in expression between control and high heteroplasmy cells (Fig.5a-b). This together with the observation that no core pluripotency genes were down-regulated in our transcriptomic analysis (Table S2a-b), suggested that the pluripotent status of mutant cells was unaffected. To further test this possibility, we differentiated the cells using monolayer assays into the neural, mesoderm and endoderm lineages. When this was done, we found that high heteroplasmy mtND6 mutant cells and controls displayed similar levels of expression of PAX6 when subjected to neural differentiation, similar levels of TBXT expression when they underwent mesoderm differentiation, and similar expression of SOX17 when induced to become endoderm (Fig.5c-d). The heteroplasmy level of the 77% mutant cells also remained stable after differentiation (Fig.S5a). These results indicate that the inherent ability of the mutant cells to undergo germ-layer differentiation is not compromised by their metabolic defects.

**Figure 5.**
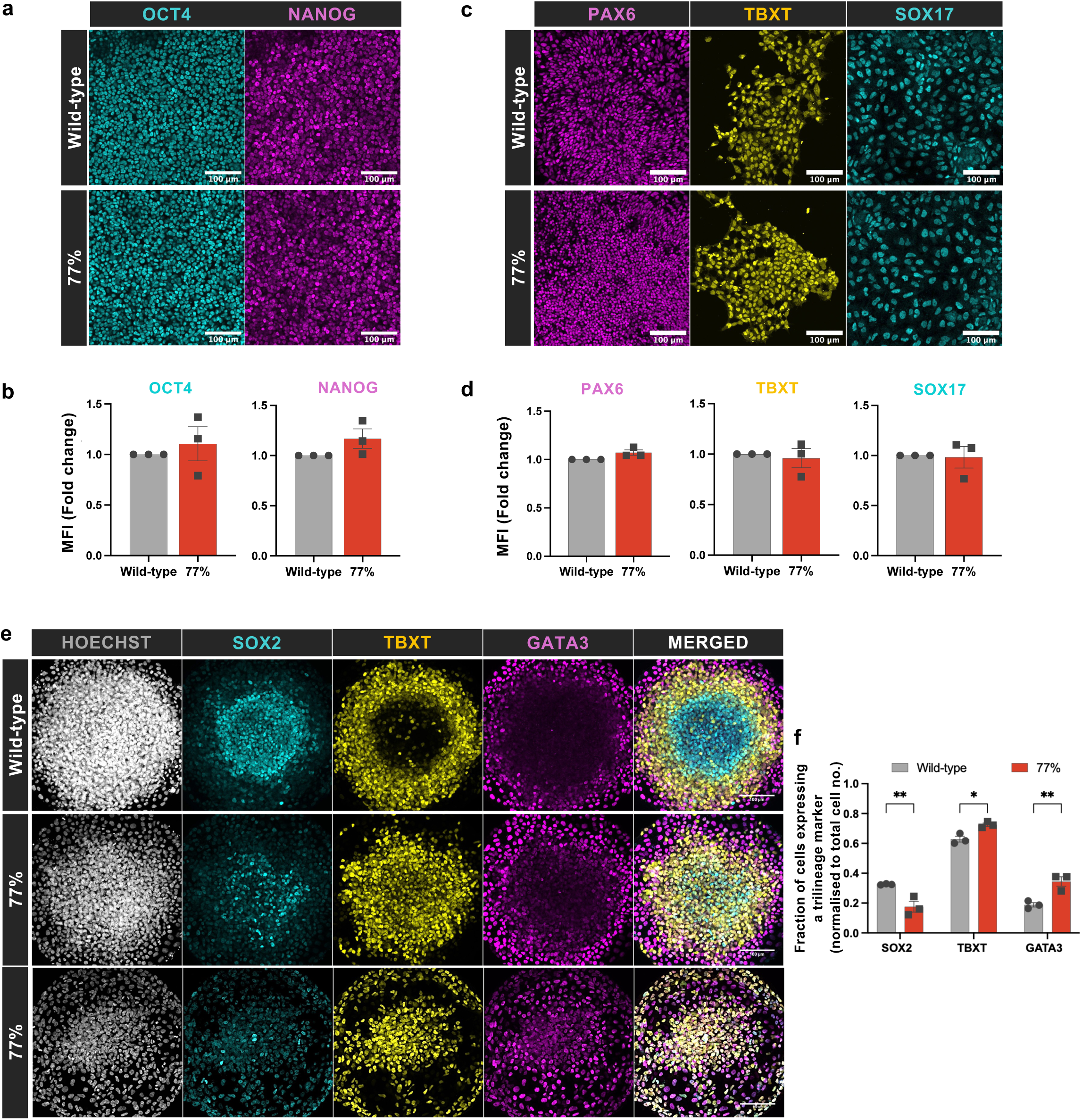
mtND6 mutation preserves pluripotency and tri-lineage differentiation capacity but impairs spatial patterning in 2D gastruloids. (a) Representative immunofluorescence images of pluripotency markers OCT4 and NANOG in wild-type and 77% mtND6 mutant hESCs cultured in mTeSR^TM^ Plus. (b) Quantification of OCT4 (left) and NANOG (right) mean fluorescence intensity (MFI) in wild-type and 77% mtND6 mutant hESCs, expressed as fold change relative to wild-type. (c) Representative immunofluorescence images of lineage-associated markers PAX6 (ectoderm), TBXT (mesoderm) and SOX17 (endoderm) in wild-type and 77% mtND6 mutant hESCs after tri-lineage differentiation. Scale bars, 100 μm. (d) Quantification of PAX6 (left), TBXT (middle) and SOX17 (right) MFI in wild-type and 77% mtND6 mutant hESCs, expressed as fold change relative to wild-type. (e) Representative immunofluorescence images of 2D gastruloids stained with Hoechst (nuclei), SOX2 (ectoderm), TBXT (mesoderm) and GATA3 (endoderm) in wild-type and 77% mtND6 mutant cells. (f) Fraction of cells expressing SOX2, GATA3, and TBXT in 2D gastruloids derived from wild-type and 77% mtND6 mutant hESCs, normalised to the total cell number. Data are shown as mean ± SEM from n=3 independent biological replicates. Statistical significance was assessed using two-tailed one-sample t-test for (b and d), or two-way ANOVA with Šídák’s post-hoc test for (f); *p < 0.05, **p < 0.01.

Embryonic development requires not only the ability to form the specific cell types that comprise the embryo, but also to organise them in space, allowing to form the structures that will give rise to the organs. This process of patterning generally involves responding in a graded fashion to the signalling factors that govern this spatial organization. We therefore wanted to analyse if mtND6 mutation affected embryonic patterning. For this we generated 2D gastruloids using micropatterns, where exogenous BMP4 induces hESCs to self-organise into an outer endodermal layer, an intermediate mesodermal layer and an inner ectodermal layer^19^. We found that in contrast to control cells that correctly spatially organised the three germ-layers, high heteroplasmy mutant cells displayed clear patterning defects. These could be most notably visualised by a reduced SOX2-positive inner ectodermal layer, that was now occupied by an expanded TBXT-positive mesodermal layer (Fig.5e, Fig.S5b). Quantification of the proportion of cells forming each germ-layer revealed that mutant cells showed an increase in the proportion of TBXT-positive mesodermal and GATA3-positive endodermal cells, that was accompanied by a decrease in the proportion of SOX2-positive ectodermal cells (Fig.5f, Fig.S5c). The decrease in SOX2-positive ectodermal cells agrees with neural differentiation being one of the GO terms identified when we compared the transcriptional profile of control and mutant hESCs (Fig.4b).

The signal that initiates germ-layer organization in these micropatterns is the TGFbeta signalling molecule BMP4^19^. We therefore tested the response of high heteroplasmy mtND6 mutant and control hESCs to this signal. Surprisingly, we observed that in mTeSR^TM^ Plus mutant cells displayed an enhanced level of phosphorylation of SMAD1/5/9, the key intracellular event that occurs upon BMP4 stimulation (Fig.6a-b). Furthermore, inhibition of Complex I activity with rotenone, or of mitochondrial translation with chloramphenicol, produced a similar increase in BMP4 signalling (Fig.6c-f). The results indicate that low Complex I activity enhances the response to BMP4.

**Figure 6.**
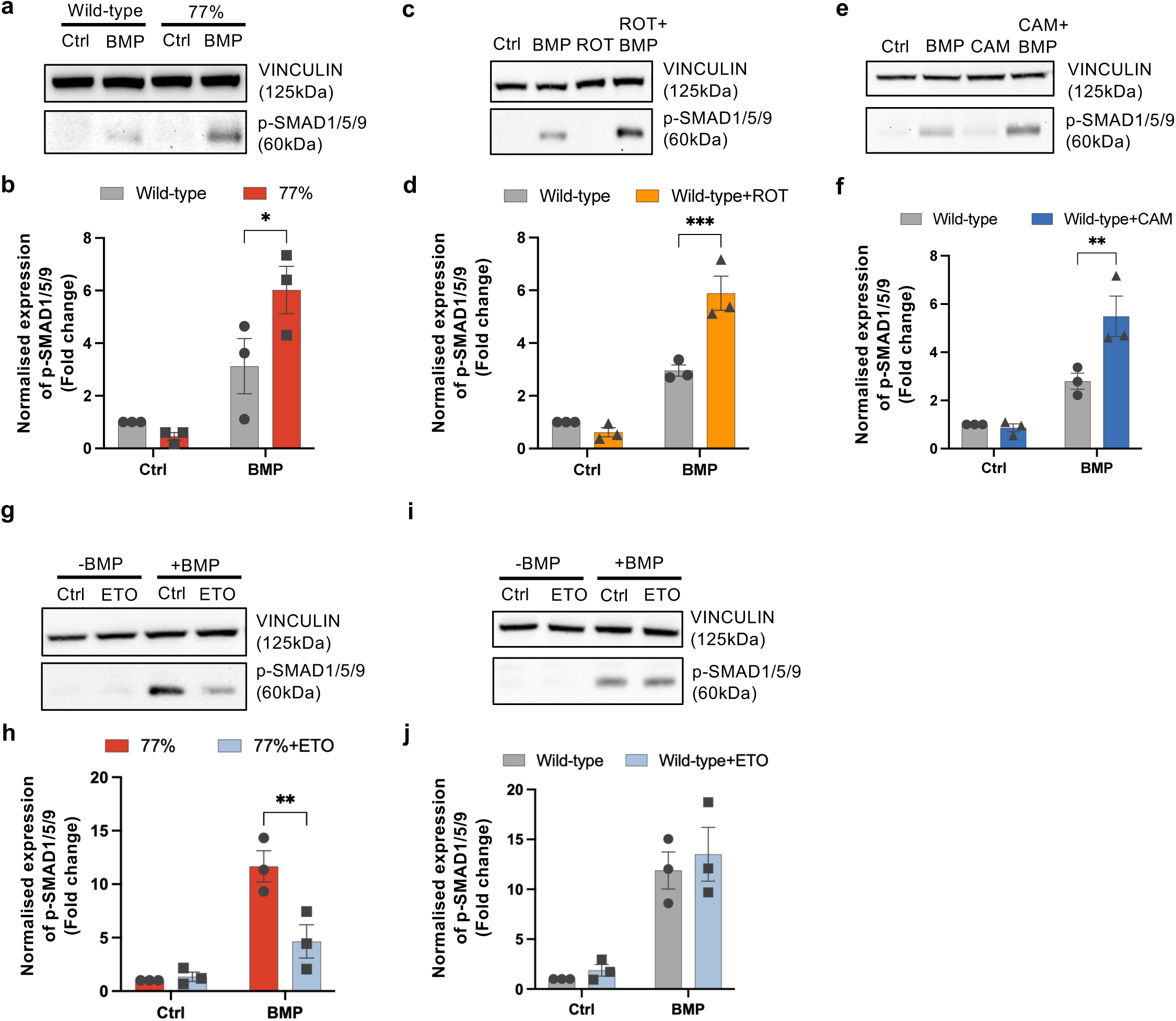
mtND6 mutation increases BMP signalling responsiveness through fatty acid oxidation-dependent mechanisms. (a) Representative immunoblot analysis of phosphorylated SMAD1/5/9 (p-SMAD1/5/9) and VINCULIN (loading control) in wild-type and 77% mtND6 mutant hESCs treated with 50ng/mL BMP or vehicle control. (b) Quantification of p-SMAD1/5/9 normalised to VINCULIN and expressed as fold change relative to control in wild-type hESCs. (c) Representative immunoblot analysis of p-SMAD1/5/9 and VINCULIN in wild-type hESCs pre-treated with 0.1μM Rotenone (ROT) for 48 hours, followed by 16 hours treatment with 50ng/mL BMP or vehicle control. (d) Quantification of p-SMAD1/5/9 levels normalised to VINCULIN and expressed as fold change relative to control in wild-type hESCs with or without Rotenone (ROT) treatment. (e) Representative immunoblot analysis of phosphorylated SMAD1/5/9 (p-SMAD1/5/9) and VINCULIN (loading control) in wild-type hESCs treated with BMP in the presence or absence of chloramphenicol (CAM). (f) Quantification of p-SMAD1/5/9 levels normalised to VINCULIN and expressed relative to control in wild-type hESCs with or without CAM treatment under control and BMP-treated conditions. (g) Representative immunoblot analysis of p-SMAD1/5/9 and VINCULIN (loading control) in 77% mtND6 mutant hESCs pre-treated with 1μM Etomoxir (ETO) for 48 hours, followed by 16 hours treatment with 50ng/mL BMP or vehicle control. (h) Quantification of p-SMAD1/5/9 normalised to VINCULIN and expressed as fold change relative to vehicle control-treated 77% mtND6 mutant hESCs. (i) Representative immunoblot analysis of p-SMAD1/5/9 and VINCULIN (loading control) in wild-type hESCs pre-treated with 1μM Etomoxir (ETO) for 48 hours, followed by 16 hours treatment with 50ng/mL BMP or vehicle control. (j) Quantification of p-SMAD1/5/9 normalised to VINCULIN and expressed as fold change relative to vehicle control-treated wild-type hESCs. Data are shown as mean ± SEM from n=3 independent biological replicates. All statistical significance was assessed using two-tailed one-sample t-test; *p < 0.05, **p < 0.01, ***p < 0.001.

Given that we had observed that mutant cells used fatty acid oxidation as a compensatory mechanism for their metabolic defects, we tested if increased beta-oxidation was causing the increase in BMP signalling. When cells were treated with 1μM etomoxir we found that this led to a decrease in pSMAD1/5/9 in mutant cells but not in controls (Fig.6g-j). This points to the cause of the change in BMP response induced by ETC dysfunction being the activation of a compensatory beta-oxidation metabolic pathway.

Together, these results suggest that during the onset of gastrulation mitochondrial activity determines the level of BMP signalling and it does so by directing which nutrients are utilised to fuel the ETC. If cells are unable to dynamically switch their nutrient utilisation as the embryonic lineages form, for example because of mtDNA mutations, this compromises their ability to spatially organise the germ layers, leading to patterning defects.

## Discussion

It is estimated that about 40% of pregnancies are lost around the time of implantation^52^ and we have previously shown that during early mouse development, mitochondrial DNA mutations are associated with patterning defects occurring around the time of gastrulation^53^. Here, we have found that mtND6 mutations that reduce OxPhos activity do not affect the ability of hESCs to differentiate into endoderm, mesoderm and ectoderm when cells are induced to take on only one of these fates. Instead, in 2D gastruloids that recapitulate the signalling relays occurring during normal development^54^, these cells are unable to self-organise into appropriately segregated mesodermal, endodermal and ectodermal domains. We show that a likely cause for this inability is increased fatty acid oxidation in mtND6 mutant cells, as this enhances their response to BMP4, the signal that initiates this self-organization. Therefore, our findings implicate embryo loss at gastrulation as a potential contributor to the selection against pathogenic mtDNA mutations occurring during development^6, 8^.

Our work also has broader implications for how patterning emerges during gastrulation and the role of mitochondrial activity. We find that mitochondrial activity plays is key for establishing the precise signalling levels that are activated in response to BMP4. Our results suggest that mitochondria do so by directing which nutrients are utilised by the ETC; when fatty acids are taken up by macropinocytosis and oxidised instead of glucose, this leads to an enhanced response to BMP4. Importantly, we show that cells need to be able to dynamically switch between the type of which nutrients they utilise, as when they can’t, for as in the mtND6 mutant cells, this causes patterning defects.

The significance that nutrients have in imparting cell fate and in patterning is only starting to be fully appreciated. Recent work has identified that glycolysis plays instructive roles in mesoderm and endoderm specification^12–14^, as well as in regulating mesoderm migration during gastrulation^12^. Interestingly, these studies found metabolites involved in hexosamine biosynthesis, rather than glucose oxidation, were responsible for these roles. During early mouse development lipid accumulation and mobilization occurs as cells change their pluripotent cell identity, and these changes drive epithelial organization during differentiation^55^. Similarly, lipid composition has been shown to affect the pluripotent state of hESCs^56^. Importantly, in hESCs fatty acid oxidation promotes endoderm differentiation by regulating the nuclear localization of SMAD3^17^. This suggests that different nutrients play different roles in different tissues, but how a variable nutrient intake, as can occur due to changes in maternal diet, can be translated into a stereotypical patterning response, is very much an open question.

Our study also has relevance to our understanding of mitochondrial DNA mutations, that are causative to mitochondrial disorders^3^, the largest group of inborn errors of metabolism, and contribute to the development of conditions as diverse as cancer^57^ and neurodegeneration^58^, as well as being associated with ageing^59^. Here, we have found that high heteroplasmy mutations in the ETC Complex I component mtND6 disrupts OxPhos leading to a compensatory change in nutrient utilisation in hESCs. We show that these cells instead of relying on glucose for their proliferation, switch to fatty acids uptake via macropinocytosis and their subsequent oxidation. However, we also demonstrate that these metabolic changes impair the ability of mutant cells to properly respond to BMP4, one of the primary signals involved in patterning the gastrulating embryo. Consequently, mtND6 mutant cells display disrupted organization of the 3 germ-layers in gastruloid models, providing a direct link between how pluripotent cells uptake nutrients and their ability to properly undergo patterning.

Changes in nutrient utilisation have been previously shown to occur as a compensatory response to mtDNA mutations. For example, m.3243 A > G mutations in the tRNA^Leu^ cause human induced pluripotent stem cells to become reliant on glycolysis^60^. Similarly, m.8993 T > G mutations cause cells to become dependent on reductive carboxylation of glutamine to sustain their proliferation^61, 62^. Our finding that hESCs with mtND6 mutations upregulate fatty acid metabolism, and require fatty acid oxidation for their proliferation, contrasts with these studies regarding the nature of the nutrients that act as alternative sources of ATP generation. They are however consistent with the overall concept that pluripotent cells have the plasticity to rewire their metabolic pathways to deal with impaired OxPhos activity. One possible explanation for this metabolic switch is that unlike during glucose oxidation, where nearly all electrons are channelled through Complex I, fatty acid oxidation produces more FADH2, that donates electrons directly to the CoQ pool, bypassing Complex I. Interestingly, a high-fat ketogenic diet has been shown to alleviate the symptoms from Leigh syndrome, MELAS and other Complex I deficiency patients^63–65^. Also, a high-fat diet has beneficial effects in mouse models of Complex I deficiency and mtDNA mutations^66–68^. These observations suggest the symptom amelioration observed could be due to a similar metabolic rewiring to the one we observe in mtND6 mutant hESCs. Future work in needed to test this hypothesis.

## Acknowledgments

We would like to thank members of the Rodriguez lab for critical discussions and helpful suggestions, and Prof. Hansong Ma at the University of Birmingham for insightful discussions. We thank Stephen Rothery for guidance and advice with confocal microscopy. We thank James Elliot, Joana De Teixeira Carrelha and Bhavik Patel from the LMS/NIHR Imperial Biomedical Research Centre Flow Cytometry Facility for support.

---

**Figure S1. Generation of mtND6 mutant hESC clones with different heteroplasmy levels and analysis of their mitochondrial activity.** (a) Sanger sequencing chromatograms of the mtND6 target region in wild-type and edited hESC clones (ND6_A6, ND6_A5, ND6_C16, and ND6_C18). (b) Heteroplasmy levels of m.14438C>T, m.14439C>T, m.14443G>A, and m.14445G>A of wild-type and mtND6 mutant clones (ND6_A6, ND6_A5, ND6_C16, and ND6_C18), quantified by amplicon sequencing. Values above each bar indicate the corresponding heteroplasmy percentage. (c) Oxygen consumption rate (OCR) profiles of wild-type and mtND6 mutant hESCs with indicated heteroplasmy levels measured using the Seahorse Mito Stress Assay. (d) Quantification of basal respiration, maximal respiration, proton leak, and ATP-linked respiration derived from OCR measurements. (e) Representative flow cytometry histograms of tetramethylrhodamine methyl ester (TMRM) fluorescence in wild-type and 77% mtND6 mutant hESCs. (f) Quantification of TMRM median fluorescence intensity (MFI) in wild-type and 77% mtND6 mutant hESCs, expressed as fold change relative to wild-type. (g) Representative flow cytometry histograms of MitoSOX fluorescence in wild-type and 77% mtND6 mutant hESCs. (h) Quantification of MitoSOX median fluorescence intensity (MFI) in wild-type and 77% mtND6 mutant hESCs, expressed as fold change relative to wild-type. Data are shown as mean ± SEM from n=3 independent biological replicates. Statistical significance was assessed using two-way ANOVA with Dunnett’s post-hoc test for (d) or two-tailed one-sample t-test for (f and h); *p < 0.05, **p < 0.01, ***p<0.001 ****p<0.0001.

**Figure S2. Characterization of mtND6 mutants and medium composition of hESCs.** (a) Proliferation rates of wild-type and mtND6 mutant hESCs (73%, and 81%) in mTeSR^TM^ Plus. (b) Comparison of the components and concentrations of E8 and mTeSR^TM^ Plus media. The table lists the concentrations of individual components formulated in each medium, with shared components shown alongside medium-specific supplements.

**Figure S3. Lipid supplementation and pharmacological targeting of lipid metabolism in wild-type and mtND6 mutant hESCs.** (a) Growth curves of wild-type and 77% mtND6 mutant hESCs cultured in E8 or E8 supplemented with chemically defined lipid concentrate (LPC). (b) Proliferation rates of wild-type and 77% mtND6 mutant hESCs in E8 or E8 supplemented with LPC. (c) Growth curves of wild-type and 77% mtND6 mutant hESCs cultured in E8 or E8 supplemented with MEM non-essential amino acid (NEAA). (d) Proliferation rates of wild-type and 77% mtND6 mutant hESCs in E8 or E8 supplemented with NEAA. (e) Growth curves of wild-type or 77% mtND6 mutant hESCs treated with 50 μM SB204990 or vehicle control. (f) Proliferation rates of wild-type hESCs following SB204990 treatment, calculated as the ratio of post-treatment to pre-treatment proliferation rates (over 48 hours). (g) Growth curves of wild-type or 77% mtND6 mutant hESCs treated with 20 μM ETC-1002 or vehicle control. (h) Proliferation rates of wild-type hESCs following ETC-1002 treatment, calculated as the ratio of post-treatment to pre-treatment proliferation rates (over 48 hours). (i) Growth curves of wild-type or 77% mtND6 mutant hESCs treated with 2 μM ACSS2i or vehicle control. (j) Proliferation rates of wild-type hESCs following ACSS2i treatment, calculated as the ratio of post-treatment to pre-treatment proliferation rates (over 48 hours). (k) Growth curves of wild-type or 77% mtND6 mutant hESCs treated with 20 μM Pravastatin or vehicle control. (l) Proliferation rates of wild-type hESCs following Pravastatin treatment, calculated as the ratio of post-treatment to pre-treatment proliferation rates (over 48 hours). Data are shown as mean ± SEM from n=3 or 4 independent biological replicates. Statistical significance was assessed using two-way ANOVA with Šídák‘s post-hoc test for (b, d, f, h, j and l); ***p<0.001, ****p<0.0001.

**Figure S4. Characterization of chloramphenicol-treated hESCs in mTeSR^TM^ Plus.** (a) Immunoblot analysis of oxidative phosphorylation (OXPHOS) complex subunits in wild-type cells treated with vehicle control or Chloramphenicol (CAM). VINCULIN was used as a loading control. (b) Quantification of NDUFB8 (Complex I), SDHB (Complex II), UQCRC2 (Complex III), MTCO1 (Complex IV), ATP5A (Complex V), protein levels, normalised to control. Data are shown as mean ± SEM from n=3 independent biological replicates. Statistical significance was assessed using two-way ANOVA with Šídák’s post-hoc test; ****p<0.0001.

**Figure S5. mtND6 mutation is maintained across differentiation and disrupts germ layer patterning in 2D gastruloids.** (a) Heteroplasmy levels of the mtDNA variant m.14438C>T in the 77% mtND6 clone measured with ARMS-qPCR in undifferentiated cells and following differentiation towards endoderm, mesoderm, and ectoderm, respectively. (b) Representative immunofluorescence images of 2D gastruloids derived from wild-type and high-heteroplasmy (81%) mtND6 mutant hESCs stained for Hoechst (nucleus), SOX2 (ectoderm), TBXT (mesoderm), and GATA3 (endoderm); n=2; scale bars, 100 μm. In (a), data are shown as mean ± SEM from n=3 independent biological replicates. Statistical significance was assessed using one-way ANOVA with Šídák’s post-hoc test.

**Figure S6. Flow cytometry gating strategy.** (a) Cell population gate: Forward scatter area (FSC-A) versus side scatter area (SSC-A) plot used to exclude debris and very large aggregates; the polygonal gate captures intact cells. (b) First doublet exclusion: FSC-H versus FSC-A gating removes doublets and multiplets, yielding a population of single cells. (c) Second doublet exclusion: SSC-H versus SSC-A gating further refines the single-cell population. (d) Live-cell gate: Sytox Blue exclusion (V450_50 channel) versus FSC-A distinguishes live (Sytoxblue-negative) cells; the boxed region indicates live single cells retained for analysis. (e) Fluorescence distribution: Histogram of YG586_15 (TMRM) fluorescence intensity, normalised to the mode, showing the mitochondrial membrane potential profile of the gated live single-cell population. Percentages of total events retained at each gating step are indicated.

## Table titles and legends

**Table S1. Mitochondrial DNA sequences in mtND6 mutant cells.** Each row represents a unique mitochondrial DNA (mtDNA) variant detected in the parental wild-type hESCs, 5%, 56% and 77% mutants in triplicates. Columns 1–4 describe the mutation: Gene indicates the mitochondrial gene or region in which the variant is located (annotated based on the revised Cambridge Reference Sequence, rCRS; NC_012920.1); Position refers to the nucleotide position on the mtDNA genome (1–16,569 bp); Wild-type and Mutation indicate the reference and observed alternative alleles, respectively. Columns 5–16 show the heteroplasmy level (%) of each variant in the corresponding sample, expressed as the percentage of reads supporting the alternative allele. Samples are grouped by experimental condition (wild-type, 5%, 56% and 77%), with three biological replicates per condition. Only variants with a heteroplasmy level ≥1% in a given sample are reported; values below this threshold are shown as blank cells to exclude low-level sequencing noise.

**Table S2 a-b. Differentially expressed genes in 77% mtND6 mutant cells compared to wild-type hESCs in mTeSR^TM^ Plus and E8.** DEGs identified by bulk RNA-sequencing between 77% mtND6 mutant and isogenic wild-type hESCs (n = 3 replicates per group) cultured in mTeSR^TM^ Plus (Table S2a) or in E8 (Table S2b). Reads were aligned to GRCh38 using STAR, gene-level counts obtained with featureCounts, and differential expression analysed with DESeq2. Columns: ID, Ensembl gene ID; log2FoldChange, log2 fold-change (77% vs. wild-type); pvalue and padj, raw and Benjamini-Hochberg-adjusted p-values; wild-type_1–3 and 77%_1–3, DESeq2-normalised counts per replicate; Gene.name, HGNC gene symbol. Significance threshold: padj < 0.05 and |log2FC| > 1.

**Table S3. Primer list for Sanger sequencing and ARMS-qPCR.**

**Table S4. Antibody list for immunofluorescence staining and western blotting.**

## Resource table

| REAGENT or RESOURCE | SOURCE | IDENTIFIER |
| --- | --- | --- |
| **Antibodies** |  |  |
| Nanog (D73G4) Monoclonal Antibody (Rabbit) | Cell Signalling Technology | Cat #4903; RRID: AB_10559205 |
| Oct-3/4 (N-19) (Mouse) | Santa Cruz | Cat #8628; RRID: AB_653551 |
| Human SOX17 Antibody (Mouse) | R&D Systems | Cat #MAB1924; RRID: AB_2195646 |
| Human/Mouse Brachyury Antibody (Goat) | R&D Systems | Cat #AF2085; RRID: AB_2200235 |
| Human/Mouse/Rat SOX2 Antibody (Goat) | R&D Systems | Cat #AF2018; RRID: AB_355110 |
| Human/Mouse GATA-3 MAb (Mouse) | R&D Systems | Cat #MAB6330; RRID: AB_10640512 |
| Anti-Brachyury / Bry antibody (Rabbit) | Abcam | Cat #ab209665; RRID: AB_2750925 |
| Goat Anti-mouse IgG (H+L) Cross-Adsorbed Secondary Antibody, Alexa Fluor™ 488 | Life Technologies | Cat #A11008; RRID: AB_143165 |
| Goat Anti-rabbit IgG (H+L) Cross-Adsorbed Secondary Antibody, Alexa Fluor™ 568 | Life Technologies | Cat #A11007; RRID: AB_141374 |
| Goat Anti-rabbit IgG (H+L) Cross-Adsorbed Secondary Antibody, Alexa Fluor™ 647 | Life Technologies | Cat #A21244; RRID: AB_2535812 |
| Anti-Vinculin Antibody (Mouse) | Sigma Aldrich | Cat #V9264; RRID: AB_10603627 |
| Total OXPHOS Rodent WB Antibody Cocktail (Mouse) | Abcam | Cat #ab110413; RRID: AB_2629281 |
| Phospho-SMAD1 (Ser463/465)/ SMAD5 (Ser463/465)/ SMAD9 (Ser465/467) Antibody (Rabbit) | Cell Signalling Technology | Cat #13820; RRID: AB_2493181 |
| Phospho-ATP-Citrate Lyase (Ser455) Antibody (Rabbit) | Cell Signalling Technology | Cat #4331; RRID: AB_2257987 |
| ATP-Citrate Lyase Antibody (Rabbit) | Cell Signalling Technology | Cat #4332; RRID: AB_2223744 |
| beta-Tubulin antibody (Rabbit) | Cell Signalling Technology | Cat #5346; RRID: AB_1950376 |
| HRP conjugated anti-mouse antibody (Horse) | Cell Signalling Technology | Cat #7076; RRID: AB_330924 |
| HRP conjugated anti-rabbit antibody (Goat) | Cell Signalling Technology | Cat #7074; RRID: AB_2099233 |
| **Chemicals, peptides, and recombinant proteins** |  |  |
| mTeSR™ Plus | STEMCELL Technologies | Cat #100-0276 |
| Geltrex™ LDEV-Free Reduced Growth Factor Basement Membrane Matrix | Gibco | Cat #A1413302 |
| ReLeSR™ | STEMCELL Technologies | Cat #100-0484 |
| STEM-CELLBANKER | Amsbio | Cat #11890 |
| TrypLE Express Enzyme | Gibco | Cat #12604-013 |
| Trypan Blue Solution, 0.4% | Gibco | Cat #15250061 |
| Y-27632 | Selleckchem | Cat #S1049 |
| DPBS | Gibco | Cat #14190250 |
| Paraformaldehyde Solution 4% in PBS | Santa Cruz | Cat #sc-281692 |
| Laminin-521 | Gibco | Cat #A29248 |
| Recombinant Human BMP-4 Protein | R&D Systems | Cat #314-BP-010 |
| Triton X-100 | Sigma Aldrich | Cat #93443 |
| Bovine Serum Albumin | Sigma Aldrich | Cat #A9418 |
| Hoechst 33342 Solution | Thermo Fisher Scientific | Cat #62249 |
| NP-40 lysis buffer | Thermo Fisher Scientific | Cat #J60766.AP |
| cOmplete™, Mini Protease Inhibitor Cocktail | Roche | Cat #11836153001 |
| RIPA buffer | Thermo Fisher Scientific | Cat #89900 |
| Phosphatase Inhibitor Cocktail | Cell Signalling Technology | Cat #5871S |
| 12% Criterion™ XT Bis-Tris Protein Gel | BioRad | Cat #3450118 |
| PVDF Transfer Membranes | Thermo Fisher Scientific | Cat #88518 |
| Skim Milk Powder | Sigma Aldrich | Cat #70166-500g |
| 10xTris-buffered Saline | BioRad | Cat #1706435 |
| Tween-20 | Sigma Aldrich | Cat #P1379-500 |
| Immobilon® Crescendo Western HRP substrate | Millipore | Cat #WBLUR0500 |
| SYTOX™ Blue Nucleic Acid Stain | Invitrogen | Cat #S11348 |
| Tetramethylrhodamine, Methyl Ester, Perchlorate (TMRM) | Invitrogen | Cat #T668 |
| MitoSOX™ Red Mitochondrial Superoxide Indicator | Invitrogen | Cat #S36008 |
| 2-NBDG | Invitrogen | Cat #N13195 |
| DQ™ Red BSA | Invitrogen | Cat #D12051 |
| Dextran, Tetramethylrhodamine, 70,000 MW, Neutral | Invitrogen | Cat #D1819 |
| Seahorse XF Calibrant Solution | Agilent | Cat #100840-000 |
| Etomoxir | MedChemExpress | Cat #HY-50202 |
| Chloramphenicol | Sigma Aldrich | Cat #C1919-5G |
| Rotenone | Sigma Aldrich | Cat #R8875-1G |
| SB204990 | Selleckchem | Cat #E1287 |
| ETC-1002 | Selleckchem | Cat #S7953 |
| ACSS2i | Selleckchem | Cat #S8588 |
| Pravastatin | MedChemExpress | Cat #HY-B0165A |
| BMP4 | R&D Systems | Cat #314-BP-010 |
| Fatty acid free BSA | Sigma Aldrich | Cat #A8806 |
| Chemically defined lipid concentrate | Gibco | Cat #11905031 |
| MEM Non-essential amino acid solution (100x) | Gibco | Cat #11140050 |
| **Critical commercial assays** |  |  |
| QIAGEN Plasmid Midi Kit | Qiagen | Cat #12191 |
| STEMdiff™ Trilineage Differentiation Kit | STEMCELL Technologies | Cat #05230 |
| hESC-compatible CYTOO® Arena 500 A micropattern chips | CYTOO | Cat #10-024-00-18 |
| DNeasy Blood & Tissue Kit | Qiagen | Cat #69504 |
| HOT FIREPol® DNA Polymerase | Solis BioDyne | Cat #01-02-KIT-00500 |
| QIAquick PCR Purification Kit | Qiagen | Cat #28104 |
| BCA protein assay | Thermo Fisher Scientific | Cat #23225 |
| Complex I activity assay | Abcam | Cat #ab109721 |
| Complex IV activity assay | Abcam | Cat #ab109909 |
| Seahorse XF Cell Mito Stress Test Kit | Agilent | Cat #103015-100 |
| Seahorse XF Glycolysis Stress Test Kit | Agilent | Cat #103020-100 |
| Seahorse XF DMEM assay medium pack | Agilent | Cat #103680-100 |
| RNeasy Mini Kit | Qiagen | Cat #74106 |
| **Deposited data** |  |  |
| RNA sequencing data | The raw data is accessible through ArrayExpress with accession number: E-MTAB-17070 | https://www.ebi.ac.uk/fg/annotare/ |
| **Experimental models: Cell lines** |  |  |
| Human: H1 hESC line | WiCell Research Institute | hES Cell Line: H1 |
| **Experimental models: Organisms/strains** |  |  |
| **Oligonucleotides** |  |  |
| Sanger sequencing primer sequences | Sigma Aldrich | Table S3 |
| ARMS-qPCR Primer Sequences | Sigma Aldrich | Table S3 |
| **Software and algorithms** |  |  |
| Image J (Fiji) | Open platform | https://fiji.sc/ |
| FlowJo | BD Biosciences | https://www.flowjo.com/ |
| GraphPad Prism | GraphPad | GraphPad Prism 10.6.1 |
| **Other** |  |  |
| Countess™ 3 Automated Cell Counter | Invitrogen | Cat #A49862 |
| Neon™ Transfection System | Thermo Fisher | Cat #MPK5000 |
| BD FACSAria™ Fusion flow cytometer | BD Biosciences | N.A. |
| NanoDrop™ 8000 spectrophotometer | Thermo Scientific | Cat #ND-8000-GL |
| QuantStudio™ 6 Flex Real-Time PCR System | Applied Biosystems™ | Cat #4485691 |
| Zeiss LSM 780 inverted confocal microscope | Zeiss | N.A. |
| iBright CL1500 imaging system | Invitrogen | N.A. |
| CLARIOstar<sup>Plus</sup> microplate reader | BMG Labtech | N.A. |
| BD FACSymphony™ A3 flow cytometer | BD Biosciences | N.A. |
| KaryoStat™+ Karyotyping Service | Thermo Fisher Scientific | Cat #A52849 |
| Seahorse XFe96 Analyzer | Agilent | N.A. |

## Methods

### Cell culture

H1 human embryonic stem cell lines were a gift from the A. Bernardo Group at Imperial College London. All hESC lines, including parental H1 and DdCBE-edited mtND6 mutant clones, were confirmed to have a normal karyotype (46, XY) by using the KaryoStat™ Assay service (Thermo Fisher Scientific) and tested negative for mycoplasma contamination prior to experimental use. hESCs were maintained feeder-free in mTeSR™ Plus medium (STEMCELL Technologies, Cat #100-0276) at 37°C with 5% CO_2_ on Geltrex™ LDEV-Free Reduced Growth Factor Basement Membrane Matrix (Gibco^TM^, Cat #A1413302) coated plates. Cells were passaged at 70-80% confluency using ReLeSR (STEMCELL Technologies, Cat #100-0484) according to the manufacturer’s instructions. Cells were cryopreserved in STEM-CELLBANKER (Amsbio, Cat #11890) using controlled-rate freezing and stored in liquid nitrogen.

### Growth curves

Cell proliferation was assessed by cell counting. Cells were dissociated with TrypLE Express (Gibco^TM^, Cat #12604-013) and viable cells were quantified using 0.4% Trypan Blue (Gibco™, Cat #15250061) on a Countess™ 3 Automated Cell Counter (Invitrogen, Cat #A49862). Cells were seeded at 60,000 cells/cm^2^ in quadruplicate in medium supplemented with 10 µM Y-27632 ROCK inhibitor (Selleckchem, Cat #S1049) for the first 24 hours. ROCK inhibitor was removed thereafter, and cell numbers were quantified every 24 hours for four consecutive days.

For drug treatment experiments, compounds were added 24 hours after removal of Y-27632 and maintained throughout the assay unless otherwise specified. The following inhibitors were used: Etomoxir (MedChemExpress, Cat #HY-50202), Chloramphenicol (Sigma Aldrich, Cat #C1919-5G), Rotenone (Sigma Aldrich, Cat #R8875-1G), SB204990 (Selleckchem, Cat #E1287), ETC-1002 (Selleckchem, Cat #S7953), ACSS2i (Selleckchem, Cat #S8588), Pravastatin (MedChemExpress, Cat #HY-B0165A). Each compound was tested at multiple concentrations as indicated in the corresponding figures, typically within the following ranges: Etomoxir (1-50 µM), Chloramphenicol (150 µM), Rotenone (0.1 µM), SB204990 (50 µM), ETC-1002 (20 µM), ACSS2i (2 µM), Pravastatin (20 µM). Cell numbers were quantified every 24 hours.

For supplementation experiments, cells were cultured in E8 medium and, 24 hours after seeding (concurrent with Y-27632 removal), the medium was supplemented with one of the following: MEM Non-essential amino acids (NEAA; Gibco, Cat # 11140050; 0.1mM), fatty acid-free bovine serum albumin (FA free-BSA; Sigma, Cat # A8806, 13.8g/L), bovine serum albumin (BSA; Sigma Aldrich, Cat # A9418, 13.8g/L), or chemically defined lipid concentrate (LPC; Gibco, Cat # 11905031, 20-440 μg/mL). Supplements were maintained throughout the assay, and cell numbers were quantified every 24 hours for four consecutive days.

For the BMP4 experiments, cells were pre-treated with 0.1 µM Rotenone or 1 µM Etomoxir for 48 hours, then with 50 ng/mL BMP4 (R&D Systems, Cat # 314-BP-010) for 16 hours with the existence of the inhibitors.

### Transfection

ND6-DdCBE-right side TALE (Addgene, Cat #157841) and ND6-DdCBE-left side TALE (Addgene, Cat #157842) plasmids used for electroporation were kindly provided by David Liu (Addgene plasmid). Plasmids were purified using a QIAGEN Plasmid Midi Kit (Qiagen, Cat #12191). Synthesised oligonucleotides encoding eGFP and tdTomato were cloned into the right- and left-side DdCBE constructs, respectively, by Azenta GENEWIZ. hESCs were dissociated with TrypLE Express Enzyme (Gibco^TM^, Cat #12604039), mixed with the ND6-DdCBE TALE plasmids, and electroporated using the Neon™ Transfection System (Thermo Fisher, Cat #MPK5000). Following electroporation, cells were cultured in mTeSR™ Plus medium supplemented with 10 µM Y-27632 to enhance post-transfection survival. At 48 hours post-transfection, cells co-expressing eGFP and tdTomato were isolated by fluorescence-activated cell sorting (FACS) into 96-well plates using BD FACSAria^TM^ Fusion flow cytometer (BD Biosciences). Clonal lines were expanded following validation of on-target editing by Sanger sequencing and used for downstream characterisation.

### Trilineage differentiation

The differentiation potential of hESCs was assessed using the STEMdiff™ Trilineage Differentiation Kit (STEMCELL Technologies, Cat #05230) according to the manufacturer’s instructions. Briefly, single-cell suspensions were plated at lineage-specific densities (2.5×10^5^ cells/cm^2^ for mesoderm; 7.5×10^5^ cells/cm^2^ for ectoderm and endoderm) and cultured in STEMdiff™ ectoderm, mesoderm, or endoderm differentiation media. Differentiation was performed for 5 days (mesoderm and endoderm) or 7 days (ectoderm) with daily medium changes. Cells were fixed in 4% paraformaldehyde (Santa Cruz, Cat #sc-281692) and analysed by immunofluorescence using lineage-specific markers (PAX6 for ectoderm, TBXT for mesoderm, and SOX17 for endoderm).

### 2D gastrulation models

Micropattern-based gastrulation assays were performed using hESC-compatible CYTOO® Arena 500A micropattern chips (CYTOO, Cat #10-024-00-18) coated with laminin-521 (Gibco^TM^, Cat #A29248). hESCs maintained in mTeSR™ Plus were dissociated into single cells, seeded onto micropatterns in mTeSR™ Plus supplemented with 10 µM Y-27632, and allowed to attach. Medium was subsequently exchanged to mTeSR™ Plus alone, followed by induction with 50 ng/mL BMP4 (R&D Systems, Cat #314-BP-010) for 48 hours. Micropattern chips were fixed in 4% paraformaldehyde and processed for immunofluorescence analysis using lineage-specific markers (SOX2 for ectoderm, TBXT for mesoderm, and GATA3 for endoderm). For each experiment, five gastruloids were analyzed per biological replicate, and experiments were performed in three independent biological replicates.

### DNA extraction and Sanger sequencing

Genomic DNA was extracted from cell pellets using the DNeasy Blood & Tissue Kit (Qiagen, Cat #69504) according to the manufacturer’s instructions, and DNA concentration was measured using a NanoDrop ND-8000 spectrophotometer (Thermo Scientific, Cat #ND-8000-GL). DdCBE-targeted regions were PCR-amplified using HOT FIREPol^®^ DNA Polymerase (Solis BioDyne, Cat #01-02-KIT-00500) in 25µL reaction containing 1ng genomic DNA template. PCR was performed with an initial denaturation at 95°C for 15 min, followed by 35 cycles of 95°C for 45s, 55°C for 45s, and 72°C for 1 min, with a final extension at 72°C for 5 min. Amplicons were verified by agarose gel electrophoresis and purified using the QIAquick PCR Purification Kit (Qiagen, Cat #28104). Purified PCR products were subjected to Sanger sequencing by Azenta Genewiz (UK). Primer sequences used for PCR are provided in Table S3.

### ARMS-qPCR

Allele-specific mutation levels were quantified by amplification refractory mutation system quantitative PCR (ARMS-qPCR). Reactions were performed in technical triplicates in 384-well plates on a QuantStudio™ 6 Flex Real-Time PCR System (Applied Biosystems™, Cat #4485691) using HOT FIREPol® DNA Polymerase and hydrolysis probes. Thermal cycling was carried out with an initial activation at 95°C for 15 min, followed by 40 cycles of denaturation at 95°C for 15s, annealing at 55°C for 20s, and extension at 72°C for 40s. Cycle threshold (Ct) values were recorded and used for downstream analyses. Detailed primer and probe sequences are provided in Table S3.

### Immunohistochemistry staining

Cells were fixed in 4% paraformaldehyde for 10 minutes at room temperature. Following fixation, cells were washed three times with PBS and permeabilised with 0.4% Triton X-100 (Sigma Aldrich, Cat #93443) in PBS (PBS-T) for 15 min at room temperature. Non-specific binding was blocked by incubation with 5% Bovine Serum Albumin (BSA) (Sigma Aldrich, Cat #A9418) in 0.4% PBS-T for 1 hour at room temperature with gentle shaking. Samples were incubated with primary antibodies diluted in blocking buffer (5% BSA in 0.4%PBS-T) overnight at 4°C. After three washes with 0.1% PBS-T, cells were incubated with species-appropriate fluorescent secondary antibodies and Hoechst nuclear stain (Thermo Fisher Scientific, Cat #62249) for 1 hour at room temperature in the dark. Following additional washes with 0.1% PBS-T, samples were mounted in antifade mounting medium and stored at 4°C until imaging.

Images were acquired using a Zeiss LSM 780 inverted confocal microscope (Zeiss) with identical acquisition settings across conditions within each experiment. At least three non-overlapping fields of view were imaged and analysed per condition per biological replicate. Image processing and quantitative analyses were performed using Fiji (ImageJ). Detailed information on primary and secondary antibodies, including sources and dilutions, is provided in Table S4.

### Western blotting

Cells were lysed in NP-40 lysis buffer (Thermo Fisher, Cat #J60766.AP) or RIPA buffer (Thermo Fisher Scientific, Cat #89900) supplemented with protease inhibitors (Roche, Cat #11836153001) and phosphatase inhibitors (Cell Signalling Technology, Cat #5871), and cleared lysates were quantified using a BCA protein assay (Thermo Fisher Scientific, Cat #23225). Equal amounts of protein (15μg) were resolved by SDS-PAGE on 12% Criterion^TM^ XT Bis-Tris precast gels (BioRad, Cat #3450118) and transferred to PVDF membranes (Thermo Fisher Scientific, Cat #88518). Membranes were blocked in 5% non-fat milk (Sigma, Cat #70166-500g) or BSA (Sigma, Cat #A9418) in Tris-buffered saline (BioRad, Cat #1706435) containing Tween-20 (Sigma, Cat #P1379-500) (TBS-T) and incubated with primary antibodies overnight at 4°C, followed by HRP-conjugated secondary antibodies. Membranes were washed with TBS-T between antibody incubations. Immunoreactive bands were detected using Immobilon® Crescendo Western HRP substrate (Merck, Cat #WBLUR0500) and imaged on an iBright CL1500 imaging system (Invitrogen). Detailed antibody information is provided in Table S4.

### Complex I and IV activity

Mitochondrial respiratory chain Complex I and Complex IV activities were measured using human enzyme activity microplate assay kits (Abcam; Complex I: Cat #ab109721; Complex IV: Cat #ab109909) according to the manufacturer’s instructions. Briefly, cells were harvested, lysed, and protein concentrations were determined by BCA assay. Normalised lysates were loaded in technical duplicates onto assay plates to allow capture of target complexes.

Complex I activity was quantified by monitoring NADH-dependent reduction of the provided reporter dye, with absorbance measured at 450 nm over time. Complex IV activity was determined by measuring the oxidation of reduced cytochrome c, detected as a decrease in absorbance at 550 nm. Kinetic measurements were acquired using a CLARIOstar^Plus^ microplate reader (BMG Labtech), and enzymatic activities were calculated from the change in absorbance over time.

### Flow cytometry

Flow cytometry was used to assess mitochondrial function, glucose uptake and macropinocytosis. Cells were dissociated into single-cell suspensions and stained with the indicated fluorescent probes prior to acquisition on a BD FACSymphony™ A3 flow cytometer (BD Biosciences). Dead cells were excluded using SYTOX™ Blue Nucleic Acid Stain (Invitrogen, Cat #S11348). A minimum of 100,000 events were recorded per sample.

For mitochondrial function, single-cell suspensions were incubated with tetramethylrhodamine methyl ester (TMRM; 10nM; Invitrogen, Cat #T668) or MitoSOX™ Red (5µM; Invitrogen, Cat #S36008) for 15 min at 37°C. Glucose uptake was assessed using the fluorescent glucose analogue 2-NBDG (100µg/mL; Invitrogen, Cat #N13195) following a 1 hour pre-incubation in glucose-free medium and a 30 minutes labelling period. Macropinocytosis was quantified using DQ™ Red BSA (20µg/mL; Invitrogen, Cat #D12051) or tetramethylrhodamine-labelled dextran (70kDa; 250µg/mL; Invitrogen, Cat #D1819) after incubation under the indicated culture conditions.

All flow cytometry data were processed and quantified using FlowJo (BD Biosciences). The gating strategy used for TMRM assay is provided in Fig.S6 as an example. Median fluorescence intensity (MFI) was quantified on live, single-cell populations and normalised as indicated.

### Seahorse assay

Mitochondrial respiration and glycolytic function were measured using a Seahorse XFe96 Analyzer (Agilent) with the Seahorse XF Cell Mito Stress Test Kit (Agilent, Cat #103015-100) and XF Glycolysis Stress Test Kit (Agilent, Cat #103020-100), according to the manufacturer’s instructions. Two days prior to the assay, cells were dissociated with TrypLE and seeded at 20,000 cells per well in Seahorse XF96 microplates in mTeSR™ Plus medium supplemented with 10µM Y-27632 ROCK inhibitor, with at least six technical replicates per condition. ROCK inhibitor was removed after 24 hours. The XFe96 sensor cartridge was hydrated overnight at 37°C in Seahorse XF Calibrant Solution (Agilent, Cat #100840-000), and the analyser was warmed overnight. Assay media and compounds were freshly prepared on the day of the assay.

For the mitochondrial stress test, Seahorse XF DMEM medium was supplemented with 1mM sodium pyruvate, 2mM L-glutamine, and 10mM glucose. Basal oxygen consumption rate (OCR) was recorded, followed by sequential injections of oligomycin (final concentration 1.5µM) to determine ATP-linked respiration, FCCP (final concentration 0.5µM) to assess maximal respiratory capacity, and rotenone/antimycin A (final concentration 0.5µM) to measure non-mitochondrial respiration. Hoechst 33342 (final concentration 2.5µM) was pre-mixed in the final injection (rotenone/antimycin A) for post-assay normalisation to cell number.

For the glycolysis stress test, Seahorse XF DMEM medium was supplemented with 2 mM L-glutamine. Basal extracellular acidification rate (ECAR) was recorded, followed by sequential injections of glucose (final concentration 10mM) to initiate glycolysis, oligomycin (final concentration 1µM) to assess glycolytic capacity, and 2-deoxy-D-glucose (2-DG; final concentration 50mM) to inhibit glycolysis and confirm assay specificity. Hoechst 33342 (final concentration 2.5µM) was included in the final injection for normalisation.

Immediately after Seahorse analysis, plates were transferred to a CLARIOstar^Plus^ microplate reader (BMG Labtech) for fluorescence-based cell number normalisation. A 15×15 scan matrix (3mm scan area per well) was acquired to account for within-well cell distribution. Hoechst fluorescence was measured using top-read fluorescence detection (excitation 355/20nm, emission 455/30nm). Fluorescence intensities were normalised to blank wells using MARS analysis software and subsequently imported into Seahorse Wave software to normalise OCR and ECAR values to cell number. Standard Seahorse parameters (including basal respiration, ATP-linked respiration, maximal respiration, spare respiratory capacity, basal glycolysis, and glycolytic capacity) were calculated as instructed.

### RNA sequencing

Total RNA was extracted using the RNeasy Mini Kit (Qiagen, Cat #74106) according to the manufacturer’s instructions. Briefly, cell pellets collected using TrypLE Express were washed twice with PBS and lysed in Buffer RLT supplemented with β-mercaptoethanol to ensure efficient cell lysis and RNA stabilisation. Lysates were homogenised by vortexing and clarified by centrifugation to remove insoluble material. An equal volume of 70% ethanol was added to the lysate prior to loading onto RNeasy spin columns. Columns were washed sequentially with Buffer RW1 and Buffer RPE, and RNA was eluted in RNase-free water. RNA concentration and purity were assessed using a NanoDrop ND-8000 spectrophotometer, and samples were stored at −80°C prior to shipment to Azenta Genewiz for strand-specific RNA sequencing.

RNA libraries were prepared using a poly(A) enrichment protocol and sequenced on an Illumina NovaSeq platform to generate 150 bp paired-end reads, with an average depth of ∼30 million reads per sample. Raw sequencing reads were subjected to quality control using FastQC and trimmed using Trimmomatic (v0.36) to remove adapter sequences and low-quality bases. Filtered reads were aligned to the human reference genome (GRCh38) using STAR (v2.5.2b). Gene-level read counts were obtained using featureCounts (Subread v1.5.2), summarised at the gene_id level based on the reference annotation. Differential gene expression analysis was performed using DESeq2 in R, with genes considered significantly differentially expressed at an adjusted p-value<0.05 and |log2 fold change|>1. Heatmaps and volcano plots were generated using the pheatmap and EnhancedVolcano packages, respectively. For functional interpretation, significantly differentially expressed genes were subjected to Gene Ontology (GO) enrichment analysis (Biological Process category) using the DAVID tool, with enriched terms considered significant at an adjusted p-value<0.05.

To estimate mtDNA heteroplasmy from transcriptomic data, the Mitochondrial HEteroplasmy AnalyzeR (MitoHEAR) pipeline ^69^ was applied to bulk RNA-seq alignments using the get_heteroplasmy function. Briefly, per-base allele counts were computed using get_raw_counts_allele, followed by filtering and estimation of allele fractions using get_heteroplasmy. For each hESC clone and passage, allele fractions at each mtDNA position were quantified individually.

### Statistics analysis

All experiments were performed using three independent biological replicates, defined as separate passages of cells processed independently under identical experimental conditions, unless otherwise stated. Data are presented as mean ± standard error of the mean (SEM).

Statistical analyses were performed using GraphPad Prism v9.0 (GraphPad Software, USA). For comparisons between two groups, paired or unpaired two-tailed Student’s t-tests assuming equal variance were used as appropriate to the experimental design, as specified in the corresponding figure legends. For fold-change analyses relative to a normalised control (e.g., fold change=1), one-sample t-tests were applied. For comparisons involving more than two groups with a single independent variable, one-way analysis of variance (ANOVA) followed by appropriate multiple comparisons tests was used. For experiments involving two independent variables (e.g., genotype and treatments), two-way ANOVA followed by appropriate multiple comparisons tests was performed, as indicated in the corresponding figure legends. Statistical significance is denoted as follows: p<0.05 (*), p<0.01 (**), p<0.001 (***), p<0.0001 (****).

## Data Availability

Data were analysed with standard programs and packages, as detailed above. Authors can confirm that all relevant data are included in the paper and/ or its supplementary information files. RNA-seq raw data and processed DEG lists are available through ArrayExpress, with accession number E-MTAB-17070.

