## Supplementary material for "Mitochondrial activity directs nutrient uptake to control early human embryonic patterning": Table S3

**Table S3. Primer list for Sanger sequencing and ARMS-qPCR.**

| Assay | Primer/Probe Name | Sequence (5′→3′) |
|---|---|---|
| Sanger Sequencing | mt-ND6-Forward | TTCACCCACAGCACCAATCC |
| | mt-ND6-Reverse | GCGGTGTGGTCGGGTGTGTTA |
| ARMS-qPCR (Consensus assay) | mt-COX1-probe | FAM-CCTGACTGGCATTGTATTAGCAAACTCAT-BHQ-1 |
| | mt-COX1-Reverse | GATAGGACATAGTGGAAGTG |
| | mt-COX1-Forward | TCATCTTTCTTTTCACCGTA |
| ARMS-qPCR (ARMS assay) | mt-ND6-probe | FAM-TTGAGGTCTTGGTGAGTGTTTTAGTGGGGT-BHQ-1 |
| | mt-ND6-Reverse | CCCATCATACTCTTTCACCC |
| | mt-ND6-Forward-229G (wild-type) | CAGCGATGGCTATTGtGG |
| | mt-ND6-Forward-229A (mutant) | CAGCGATGGCTATTGtGA |
| | mt-ND6-Forward-235TCC (wild-type) | GATGGCTATTGAGRAGTATCC |
| | mt-ND6-Forward-235TTT (mutant) | GATGGCTATTGAGRAGTATTT |

Intentional mismatch (lower-case letters) was added to the antepenultimate position in the forward primers targeting both wild-type and mutant DNA. Single nucleotide polymorphisms (SNPs) - specific bases are presented as underlined letters. R represents a nucleotide wobble R=A+G.
