## Supplementary material for "Mitochondrial activity directs nutrient uptake to control early human embryonic patterning": Table S4

**Table S4. Antibody list for immunofluorescence staining and western blotting.**

**For immunofluorescence staining:**

| Antibody | Host Species | Supplier and catalogue | Dilution |
|---|---|---|---|
| NANOG | Rabbit | CST, Cat #4903T | 1:200 |
| OCT4 | Mouse | Santa Cruz, Cat #8628 | 1:500 |
| SOX17 | Mouse | R&D Systems, Cat #MAB1924 | 1:50 |
| Brachyury (TBXT) | Goat | R&D systems, Cat #AF2085 | 1:20 |
| SOX2 | Goat | R&D systems, Cat #AF2018 | 1:20 |
| GATA3 | Mouse | R&D systems, Cat #MAB6330 | 1:20 |
| Brachyury (TBXT) | Rabbit | Abcam, Cat #ab209665 | 1:1000 |
| Anti-mouse IgG (H+L) Cross-Adsorbed Secondary Antibody, Alexa Fluor™ 488 | Goat | Life Technologies, Cat #A11017 | 1:600 |
| Anti-rabbit IgG (H+L) Cross-Adsorbed Secondary Antibody, Alexa Fluor™ 568 | Goat | Life Technologies, Cat #A11011 | 1:600 |
| Anti-goat IgG (H+L) Cross-Adsorbed Secondary Antibody, Alexa Fluor™ 647 | Rabbit | Life Technologies, Cat #A21446 | 1:600 |

**For western blotting:**

| Antibody | Host Species | Supplier and catalogue | Dilution |
|---|---|---|---|
| VINCULIN | Mouse | Sigma, Cat #V9624 | 1:10,000 |
| Total OXPHOS Rodent WB Antibody Cocktail | Mouse | Abcam, Cat #ab110413 | 1:250 |
| p-SMAD1/5/9 | Rabbit | CST, Cat #13820 | 1:1000 |
| HRP conjugated anti-mouse | Goat | CST, Cat #7076 | 1:3000 |
| HRP conjugated anti-rabbit | Goat | CST, Cat #7074 | 1:3000 |
