## Supplementary material for "Mitochondrial activity directs nutrient uptake to control early human embryonic patterning": Figure S1-S6

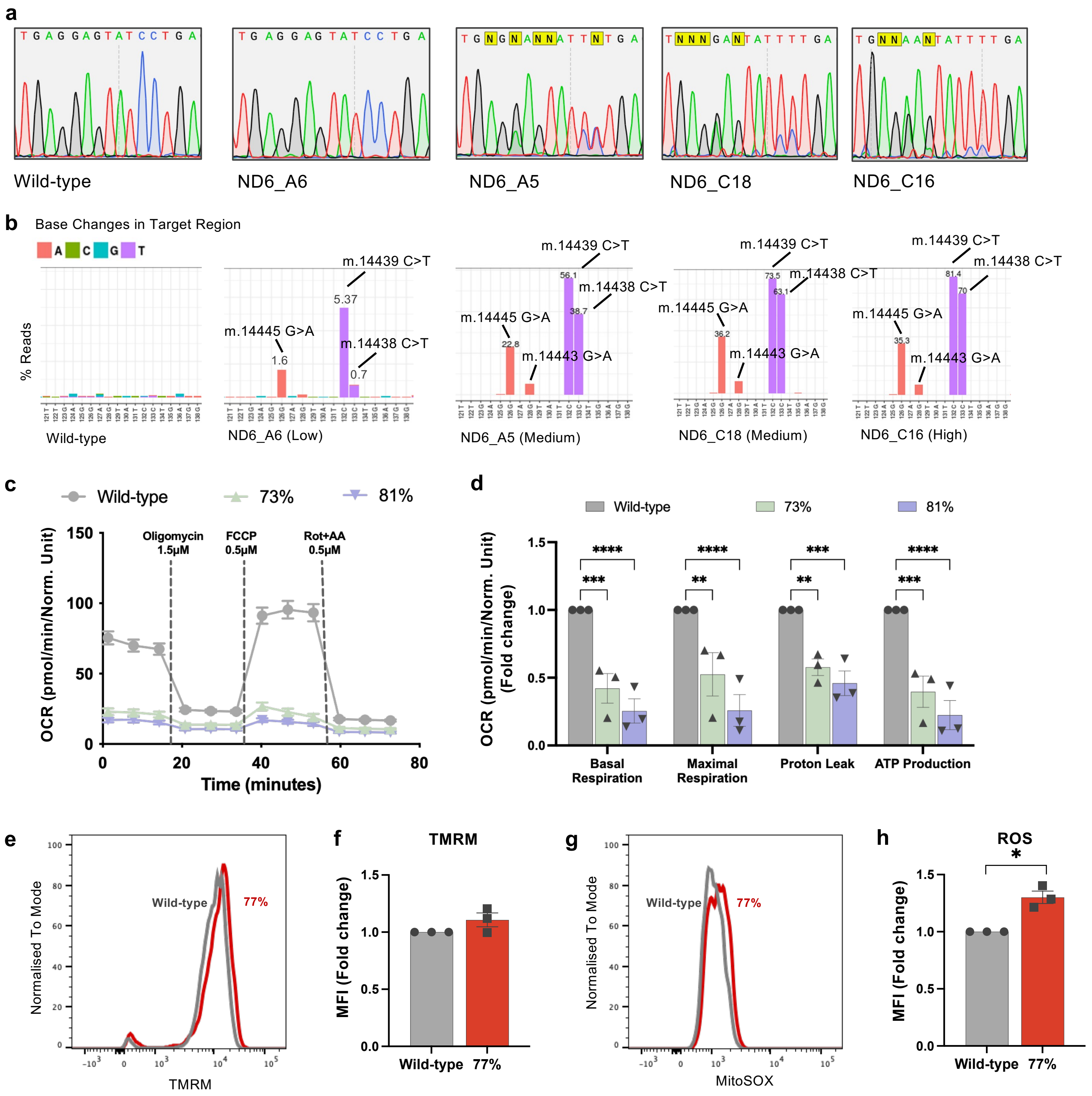

a
Wild-type
ND6_A6
ND6_A5
ND6_C18
ND6_C16
b
Base Changes in Target Region
% Reads
m.14439 C>T
m.14438 C>T
m.14445 G>A
m.14443 G>A
Wild-type
ND6_A6 (Low)
ND6_A5 (Medium)
ND6_C18 (Medium)
ND6_C16 (High)
c
Wild-type
73%
81%
Oligomycin 1.5μM
FCCP 0.5μM
Rot+AA 0.5μM
OCR (pmol/min/Norm. Unit)
Time (minutes)
d
OCR (pmol/min/Norm. Unit) (Fold change)
Basal Respiration
Maximal Respiration
Proton Leak
ATP Production
e
Normalised To Mode
Wild-type
77%
TMRM
f
TMRM
MFI (Fold change)
Wild-type
77%
g
Normalised To Mode
Wild-type
77%
MitoSOX
h
ROS
MFI (Fold change)
Wild-type
77%

### Figure S2

**a**

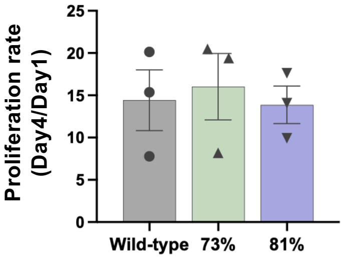


**b**

| | E8 | mTeSR+ |
|---|---|---|
| DMEM/F12 | 1000 mL | 1000 mL |
| Glucose | 17.5 mM | 17.5 mM |
| L-ascorbic Acid | 64 mg/L | 66 mg/L |
| Sodium Selenium | 14 μg/L | 14 μg/L |
| Insulin | 19.4 mg/L | 20 mg/L |
| NaHCO3 | 1383 mg/L | 560 mg/L |
| Transferrin | 10.7 mg/L | 11 mg/L |
| L-Glutamine | 292.3 mg/L | 146.15 mg/L |
| Stable FGF2 | 40 μg/L | 100 μg/L |
| TGFB1+ | 2 μg/L | 0.6 ug/L |
| BSA | / | 13.8 g/L |
| 2-Mercaptoethanol | / | 7 μL/L |
| Thiamine | / | 6.6 mg/L |
| Reduced Glutathione | / | 2 mg/L |
| Pipecolic Acid | / | 130 μg/L |
| LiCl | / | 42.36 mg/L |
| GABA | / | 103.1 mg/L |
| MEM NEAA | / | 0.1 mM |
| Chemically Defined Lipid Concentrate | / | 20-440 μg/mL |
| Trace Elements B | / | 0.24-280 μg/mL |
| Trace Elements C | / | 0.12-4.2 μg/mL |

### Figure S3

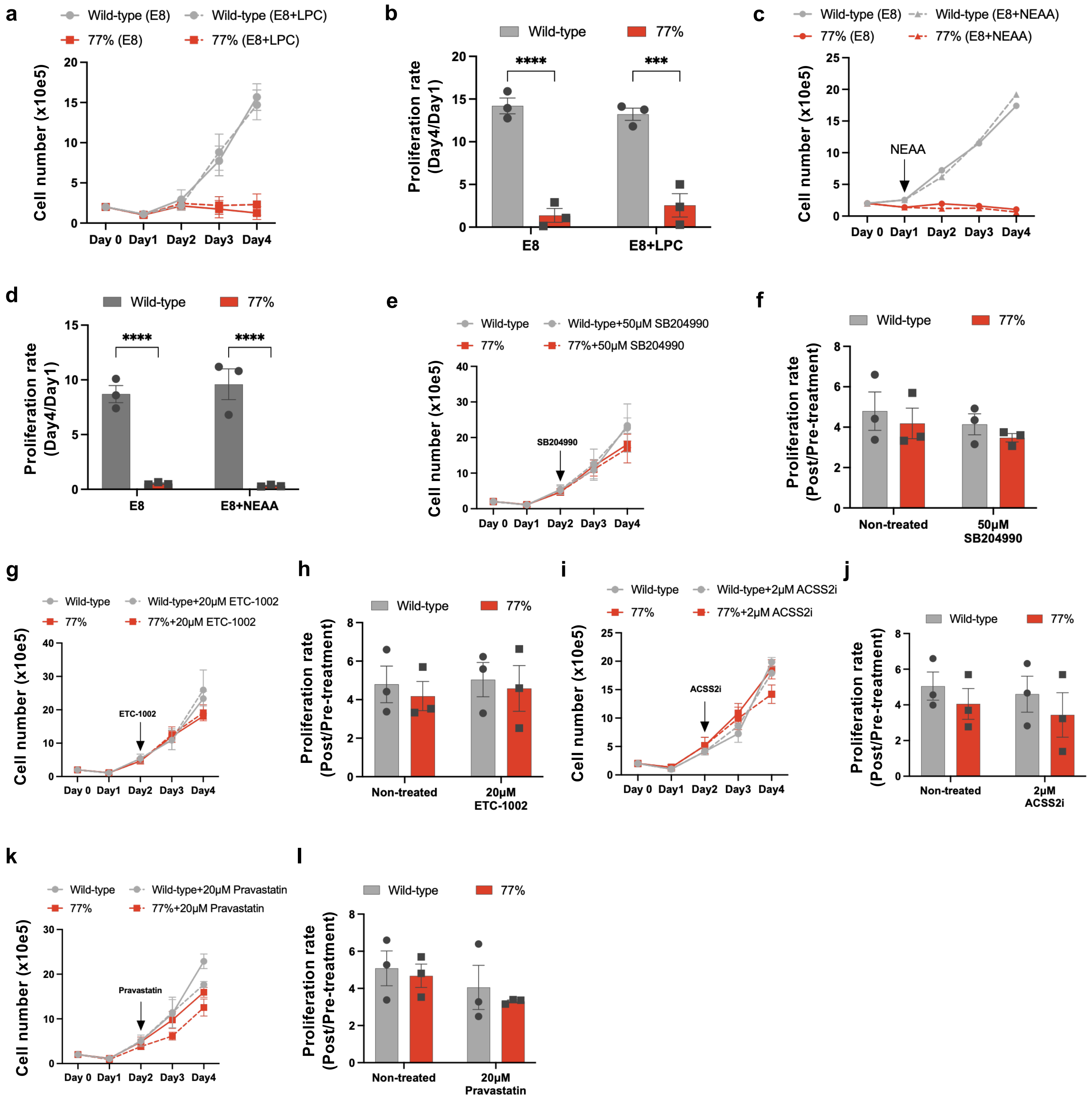

a
Wild-type (E8)
Wild-type (E8+LPC)
77% (E8)
77% (E8+LPC)
Cell number (x10e5)
Day 0 Day1 Day2 Day3 Day4
b
Wild-type
77%
Proliferation rate (Day4/Day1)
****
***
E8
E8+LPC
c
Wild-type (E8)
Wild-type (E8+NEAA)
77% (E8)
77% (E8+NEAA)
NEAA
Cell number (x10e5)
Day 0 Day1 Day2 Day3 Day4
d
Wild-type
77%
Proliferation rate (Day4/Day1)
****
****
E8
E8+NEAA
e
Wild-type
Wild-type+50μM SB204990
77%
77%+50μM SB204990
SB204990
Cell number (x10e5)
Day 0 Day1 Day2 Day3 Day4
f
Wild-type
77%
Proliferation rate (Post/Pre-treatment)
Non-treated
50μM SB204990
g
Wild-type
Wild-type+20μM ETC-1002
77%
77%+20μM ETC-1002
ETC-1002
Cell number (x10e5)
Day 0 Day1 Day2 Day3 Day4
h
Wild-type
77%
Proliferation rate (Post/Pre-treatment)
Non-treated
20μM ETC-1002
i
Wild-type
Wild-type+2μM ACSS2i
77%
77%+2μM ACSS2i
ACSS2i
Cell number (x10e5)
Day 0 Day1 Day2 Day3 Day4
j
Wild-type
77%
Proliferation rate (Post/Pre-treatment)
Non-treated
2μM ACSS2i
k
Wild-type
Wild-type+20μM Pravastatin
77%
77%+20μM Pravastatin
Pravastatin
Cell number (x10e5)
Day 0 Day1 Day2 Day3 Day4
l
Wild-type
77%
Proliferation rate (Post/Pre-treatment)
Non-treated
20μM Pravastatin

### Figure S4

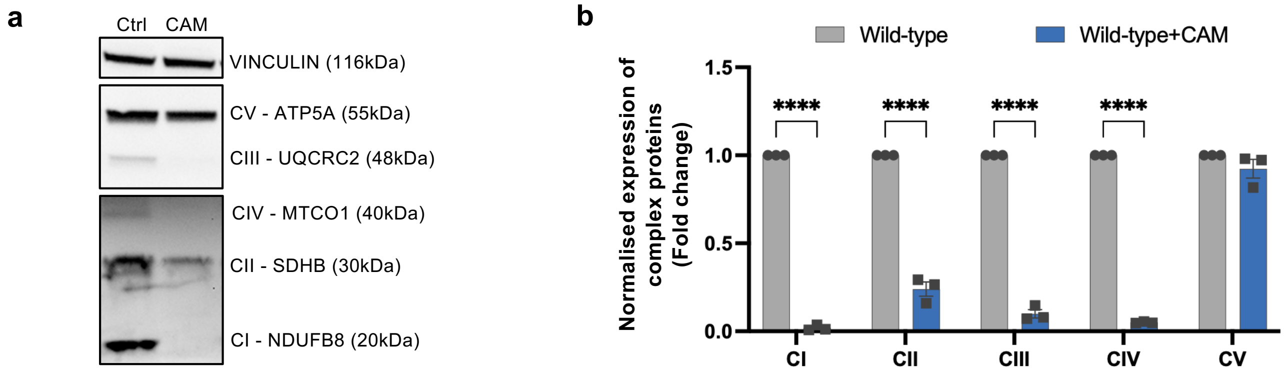

a
Ctrl
CAM
VINCULIN (116kDa)
CV - ATP5A (55kDa)
CIII - UQCRC2 (48kDa)
CIV - MTCO1 (40kDa)
CII - SDHB (30kDa)
CI - NDUFB8 (20kDa)
b
Wild-type
Wild-type+CAM
Normalised expression of complex proteins (Fold change)
1.5
1.0
0.5
0.0
****
****
****
****
CI
CII
CIII
CIV
CV

### Figure S5

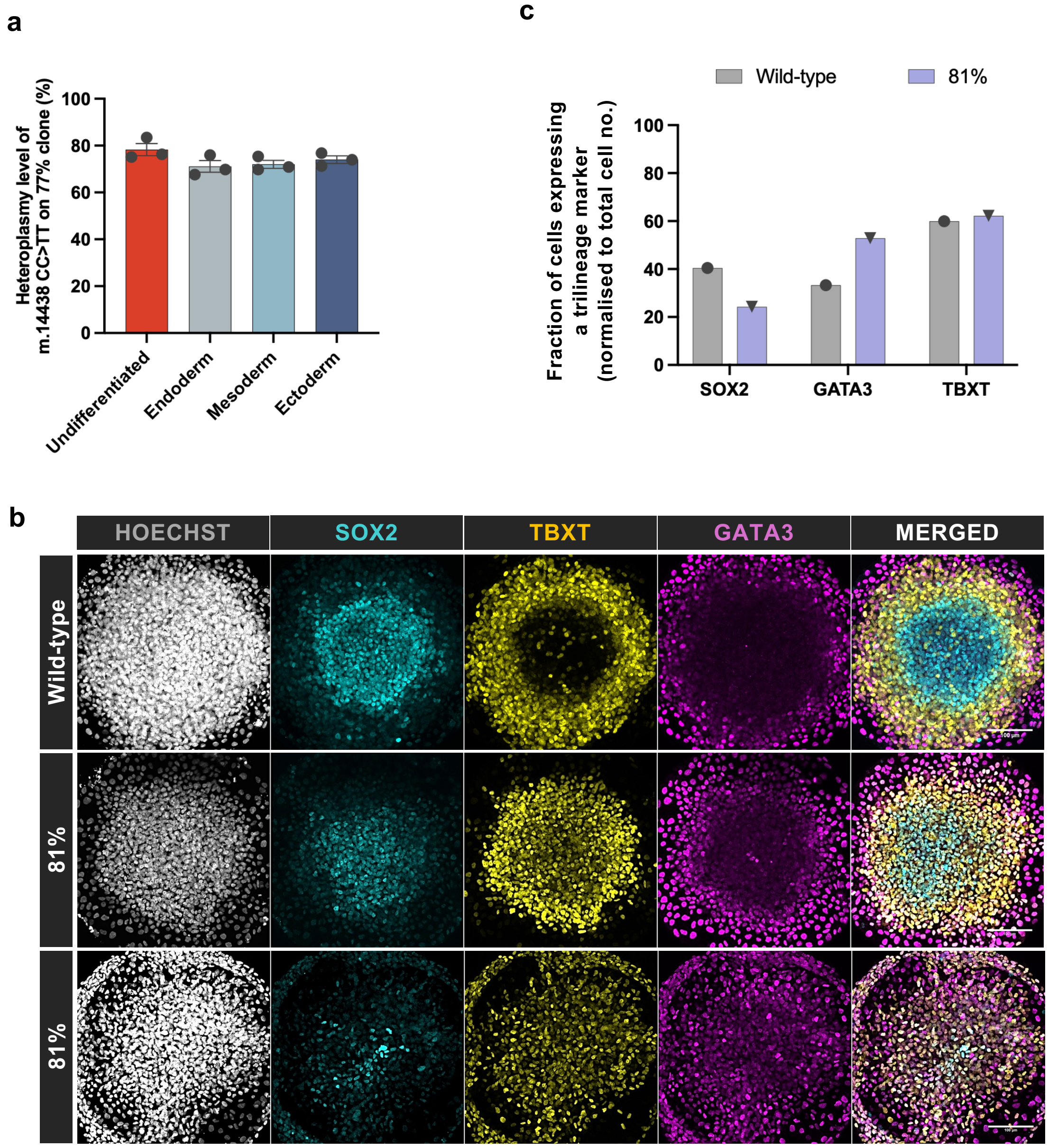
a
Heteroplasmy level of m.14438 CC>TT on 77% clone (%)
100
80
60
40
20
0
Undifferentiated
Endoderm
Mesoderm
Ectoderm
c
Wild-type
81%
Fraction of cells expressing a trilineage marker (normalised to total cell no.)
100
80
60
40
20
0
SOX2
GATA3
TBXT
b
HOECHST
SOX2
TBXT
GATA3
MERGED
Wild-type
81%
81%

### Figure S6

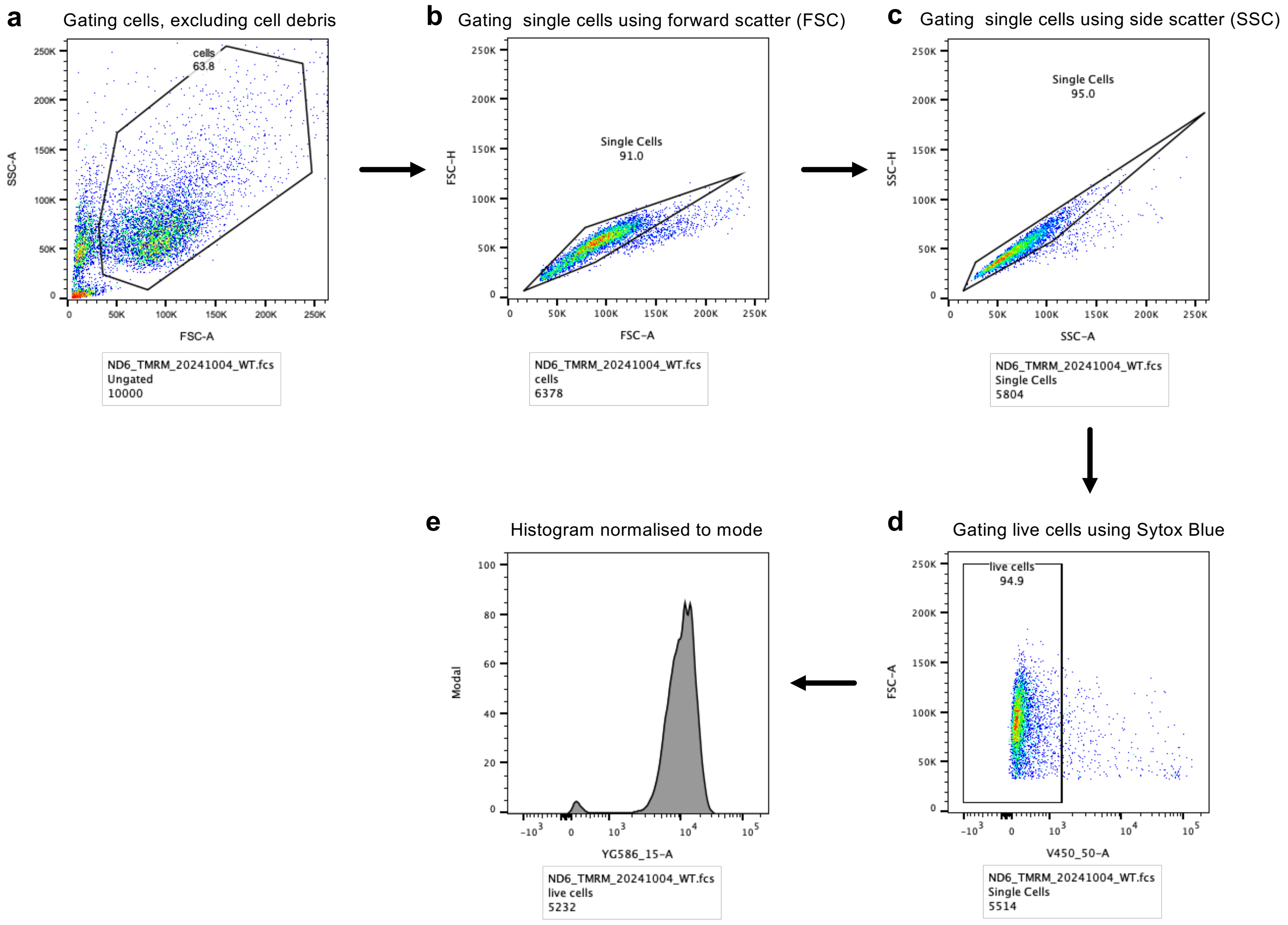
a
Gating cells, excluding cell debris
cells
63.8
SSC-A
FSC-A
ND6_TMRM_20241004_WT.fcs
Ungated
10000
b
Gating single cells using forward scatter (FSC)
Single Cells
91.0
FSC-H
FSC-A
ND6_TMRM_20241004_WT.fcs
cells
6378
c
Gating single cells using side scatter (SSC)
Single Cells
95.0
SSC-H
SSC-A
ND6_TMRM_20241004_WT.fcs
Single Cells
5804
d
Gating live cells using Sytox Blue
live cells
94.9
FSC-A
V450_50-A
ND6_TMRM_20241004_WT.fcs
Single Cells
5514
e
Histogram normalised to mode
Modal
YG586_15-A
ND6_TMRM_20241004_WT.fcs
live cells
5232
